# Structural determinants of pure antiestrogenicity

**DOI:** 10.1101/2023.10.31.564979

**Authors:** M. Diennet, F. Shaikh, M. El Ezzy, K. Thiombane, S. Weber, J. Poupart, R. Mendoza-Sanchez, Z. Gao, J. Gleason, D.A. Schuetz, G. Greene, A. Marinier, S.W. Fanning, S Mader

## Abstract

Inhibitory activities of antiestrogens on estrogen receptor alpha (ERα) range from mixed antagonism/agonism (selective ER modulators; SERMs) to complete antiestrogenicity associated with accelerated ERα turnover (selective ER degraders; SERDs). Here, we show using a panel of SERMs, SERDs and a PROTAC that efficient induction of ERα SUMOylation better accounts for complete transcriptional repression than increased suppression of coactivator recruitment or accelerated ERα degradation. Antiestrogen-induced ERα SUMOylation depends on the hydrophobicity of N-terminal residues of ligand binding domain (LBD) helix 12 (H12). L536 mutations, including those occurring in endocrine therapy-resistant breast cancer, abolished SUMOylation with all antiestrogens. Structures of the L536S human ERα LBD bound to fulvestrant analogs and molecular dynamics simulations predict dynamic side chain interactions with wild-type ERα H12 in the coactivator-binding groove and model the impact of L536 mutations. Together, our results show that induced SUMOylation contributes to the complete suppression of ERα activity and provide insights into its structural determinants.

## Main

Antiestrogens (AEs), widely used in breast cancer therapy, inhibit estrogen receptor alpha (ERα), which drives cell proliferation in up to 70% of tumors ^1–3^. ERα, a nuclear receptor and ligand-dependent transcription factor, is composed of a central DNA binding domain (DBD) mediating binding to estrogen response elements (EREs), an N-terminal activation function 1 (AF1) and a C-terminal AF2 that overlaps the ligand binding domain (LBD) ^4–8^. ERα agonists stabilize the LBD C-terminal helix 12 (H12) on top of the ligand binding pocket, creating a hydrophobic groove recognized by coactivators bearing hydrophobic α-helical LXXLL motifs ^3,9–11^. AEs contain a steroidal or non-steroidal core flanked by an extended aliphatic side chain of variable length and nature that blocks adoption of an agonist conformation by H12. Tamoxifen, the gold standard AE for treatment of primary breast tumors, is a selective estrogen receptor modulator (SERM) with partial agonist activity in uterus and bone ^2,3,12,13^. Fulvestrant (ICI182,780), dubbed a “pure” antiestrogen, lacks agonist activity ^14^. Unlike tamoxifen, it induces ERα proteasomal degradation ^15^, hence the name selective estrogen receptor degrader/downregulator (SERD) ^2,3,13^. Fulvestrant, with its long hydrophobic side chain, is poorly orally bioavailable ^16,17^. Novel SERDs with increased oral bioavailability have shorter side chains containing basic amines (e.g. RAD1901, Elacestrant; GDC0927; GDC9545, Giredestrant; AZD9833, Camizestrant) or terminating with acidic groups (e.g., AZD9496, GW5638/GW7604, GDC0810) ^3,11,18,19^. Alternatively, PROTACs (proteolysis targeting chimeras; e.g. ARV471) induce ERα degradation by direct recruitment of an ubiquitin E3 ligase ^19^.

Whether induction of ERα degradation contributes to AE therapeutic efficacy or lack of tissue-specific agonism of SERDs is currently debated ^20–24^. Notably, we found that fulvestrant and long side chain analogs induce ERα SUMOylation, contributing to efficacious transcriptional repression even in the absence of induced degradation ^23,25,26^. We report here that in a diverse panel of AEs inducing varying degrees of ERα SUMOylation, efficient SUMOylation correlates better with complete suppression of ERα transactivation than suppressed coactivator recruitment or induction of receptor degradation. To remedy the lack of crystal structures of fulvestrant or long hydrophobic side chain analogs with ERα, we solved the structures of fulvestrant analogs complexed to a L536S ERα mutant, used previously to obtain crystals with shorter side chain SERDs ^27–29^, and performed molecular dynamics simulations of fulvestrant and analogs bound to the LBD of WT ERα or of the L536R mutant, associated with clinical hormonotherapy resistance^30^. We find that induction of ER SUMOylation is an integral component of pure antiestrogenicity and is associated with the capacity of long AE side chains to interact with H12 positioned in the ERα coactivator binding groove. These studies provide a structural and mechanistic basis for pure antiestrogenicity and should inform further development and use of pure AEs.

## RESULTS

### AEs with long aliphatic side chains have increased efficacy of transcriptional suppression

To compare efficacy of different AEs, we characterized suppression of ERα transcriptional activity using an (ERE)3-TATA-Luc reporter assay in HEK293 cells treated with 4-hydroxytamoxifen (4-OHT, the active derivative of Tam), or with AEs containing side chains of varying lengths and composition (**Supplementary Fig.1)**. These included SERM/SERDs Raloxifene (Ral), Lasofoxifene (Las), Basedoxifene (Baz), novel generation SERDs GDC0927, Giredestrant (Gir), AZD9833, and long side chain “pure” AEs ICI164,384 and fulvestrant (Fulv), as well as a derivative with a side chain shorter by 1 carbon, desmethyl-ICI164,384 (UM0148682). We also tested acidic side chain SERDs (AZD9496, GDC0810, GW7604) and PROTAC (ARV471). While 17β-estradiol (E2, 10 nM) promoted ERα transactivation, 4-OHT (100 nM) had no significant effect relative to vehicle (**Fig. 1a**). All other AEs (100 nM) significantly repressed basal reporter activity (inverse agonism), with fulvestrant being the most efficacious (**Fig. 1a**). Acidic side chain AEs had only modest inverse agonist activity compared to SERDs with long aliphatic side chains, and PROTAC ARV471 was entirely devoid of inverse agonism (**Supplementary Fig. 2a**). Similar results were obtained with 1 μM AEs (**Supplementary Fig. 2b**). In osteosarcoma U2OS cells, in which OHT has partial agonist activity, inverse agonism was observed with most other AEs, with long aliphatic chains being the most efficacious, while ARV471 was a weak partial agonist (**Supplementary Fig. 3a**). We also tested repression of transcription driven by VP16-ERα, a chimera containing a strong constitutive activation function. Only AEs with the longest hydrophobic side chains (Gir, AZD9833, C10(15), ICI164,384 and fulvestrant, **Supplementary Fig. 1**) significantly repressed VP16-ERα (**Supplementary Fig. 3c-d**), confirming their superior AE activities.

**Figure 1:**
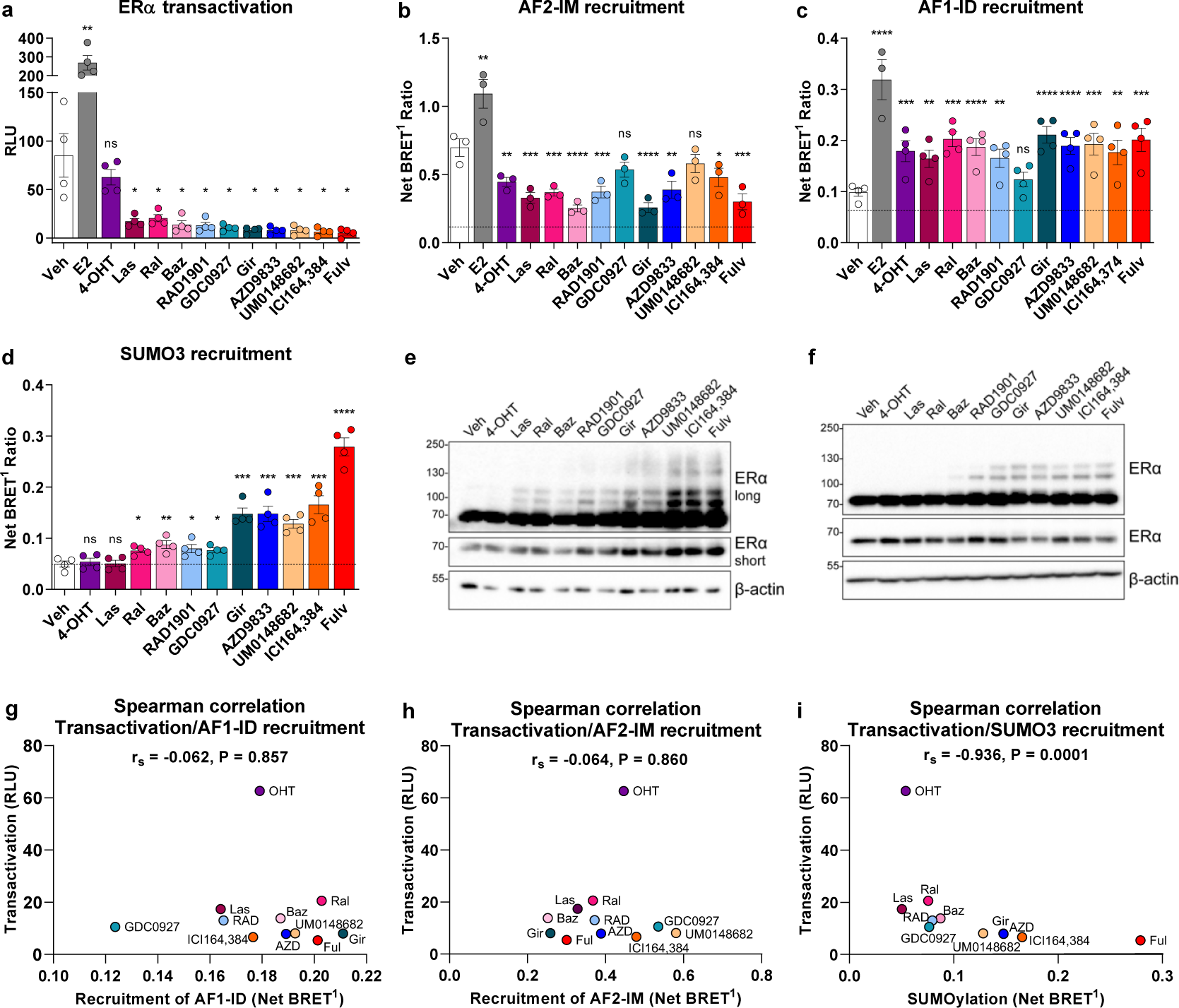
Suppression of ERα transactivation by antiestrogens correlates with receptor SUMOylation. (**a**) Differential transcriptional repression by SERMs and SERDs an ERE3-TATA-LUC reporter vector in transiently transfected HEK293 cells. Cells maintained in estrogen-depleted medium were treated with estradiol (E2, 10 nM) or AEs (100 nM) for 24 h relative luciferase values are shown (N=3, mean values +/- SEM). (**b**,**c**) Impact of AEs on recruitment of coactivator motifs/domains in BRET assays with AF2IM-YFP (b) or AF1ID-YFP (c) in transiently transfected HEK293 cells treated with E2 (10 nM) or AEs (100 nM) for 45 min. (N=3, mean values +/- SEM; dashed line: background BRET with the dimerization-defective L507R ERα mutant). (**a**,**b**,**c**) Statistical significance in a Holm-Sidak multiple comparison test is shown for the comparison of treatment vs vehicle (Veh) * P ≤ 0.05, ** P ≤ 0,01; *** P ≤ 0,001; **** P ≤ 0,0001; “ns” P > 0,05). (**d**) Impact of AEs on ERα-SUMO3 interaction in a BRET assay. (N=3, mean values +/- SEM; dashed line: net BRET1 value of the vehicle (Veh) control). (**e**,**f**) Western analysis of whole cell extracts from transfected HEK293 cells (e) or ER+ MCF-7 cells (f) cultured in estrogen-depleted media and treated for 1h with 100 nM AEs. Two different exposures are displayed to reveal protein levels of unmodified ERα (short exposure) and ligand-induced post translational modifications (long exposure). N=3, one representative blot shown. (**g**,**i**) Correlation between ERα transactivation levels and the recruitment of SUMO3 (g), AF2-IM (H) or AF1-IF (i) in a Spearman’s rank correlation analysis. Correlation coefficient (r_s_) and p value (P) are indicated on each graph.

### AE antagonism does not correlate with reduced coactivator recruitment to AF1

The gene- and tissue-specific partial agonist activity of SERMs like tamoxifen has been ascribed to the context-dependent AF1 activity of DNA-bound ERα ^31–34^. To determine whether varying residual activities of AEs result from differential recruitment of cofactors by AF1 or AF2 ^35,36^, we used BRET assays to monitor interactions of ERα with an AF2-interaction motif (AF2IM) or an AF1-interaction domain (AF1ID) (see Material and Methods). As expected, E2 (10 nM) induced coactivator domain recruitment to AF1 and AF2, in spite of the elevated concentrations of these domains driving basal recruitment (**Fig. 1b,c**). Net BRET signals in the absence of treatment were abolished by an ERα dimerization mutation (L507R) ^35^ (**Fig. 1b,c**, dashed line), indicating background activity. 4-OHT significantly reduced recruitment to AF2, but, as previously reported ^32^, induced partial recruitment to AF-1 (**Fig. 1b,c**). Other AEs variably repressed recruitment of AF2; Baz, Gir and Fulv were the most efficacious, while GDC0927 and UM0148682 lacked efficacy at 100 nM. All antiestrogens except GDC0927 induced comparable recruitment to AF1 at 100 nM. Similar results were obtained with the Receptor Interacting Domain of co-activator SRC1 as with the isolated interacting motif (**Supplementary Fig. 4**). Thus, maximal suppression of ERα transcriptional activity in reporter assays does not result from greater suppression of ERα coactivator recruitment, suggesting other mechanism(s) for the differential transcriptional repression of AEs in this panel.

### Inverse agonism correlates with induced SUMOylation rather than SERD activity

As ERα SUMOylation is important for transcriptional repression by fulvestrant ^23,25^, we examined whether repression of ERα transcriptional activity by our AE panel correlates with induced SUMOylation. AEs treatment resulted in differing degrees of SUMO3 recruitment to ERα in BRET assays, with the strongest signals observed with those with SERD activity (fulvestrant or analogs ICI164,384 and UM0148682 and new generation SERDs AZD9833 and Gir; **Fig. 1d**). Conversely, SERDs with acidic side chains did not elicit significant recruitment in HEK293 cells (**Supplementary Fig. 2e**). SUMO BRET signals correlated well with modifications of exogenous ERα in transfected HEK293 (**Fig. 1e and Supplementary Fig. 2c,** ladder of higher molecular weight bands) or U2OS cells (**Supplementary Fig. 3b**), and of the endogenous receptor in MCF-7 ER+ breast cancer cells (**Fig. 1f**). The identity of the bands as SUMOylated ERα forms was confirmed by their dose-dependent suppression by the SENP1 deSUMOylase (**Supplementary Fig. 2d**). Importantly, SENP1 coexpression in reporter assays derepressed activities of all AEs to levels similar to those of OHT (**Supplementary Fig. 2a,b**). Quantitation of residual transcriptional activity with our AE panel indicated high anticorrelation with SUMO3 recruitment (r<-0.90, Spearman correlation, **Fig. 1g**; **Supplementary Fig. 2e,f**), but lack of correlation with the level of coactivator recruitment to AF1 or AF2 (**Fig. 1h,i**), and poor correlation with endogenous ERα degradation in MCF7 cells (**Supplementary Fig. 2g,h**), consistent with a general role of SUMOylation in transrepression by a variety of AEs.

### Residues near and within H12 are key for fulvestrant-induced SUMOylation

Mutations of several bulky hydrophobic amino acids within the loop connecting H11 and H12 (H11-H12 loop) or in H12 increase the agonist activity of antiestrogens ^20,22,25,37^. To determine their contributions to induced ERα SUMOylation, we performed an alanine-scanning mutagenesis of the H11-H12 loop and H12. Substitution of bulky hydrophobic residues at positions 536 and 541 led to constitutive ERα activity in HEK293 and U2OS cells (**Fig. 2a**, **Supplementary Fig. 5a**), as reported previously ^20,38^. Conversely, H12 mutants L539A, L540A and M543A had decreased basal activity (**Fig. 2a**), correlating with reduced AF2IM recruitment in BRET assays (**Supplementary Fig. 6**).

**Figure 2:**
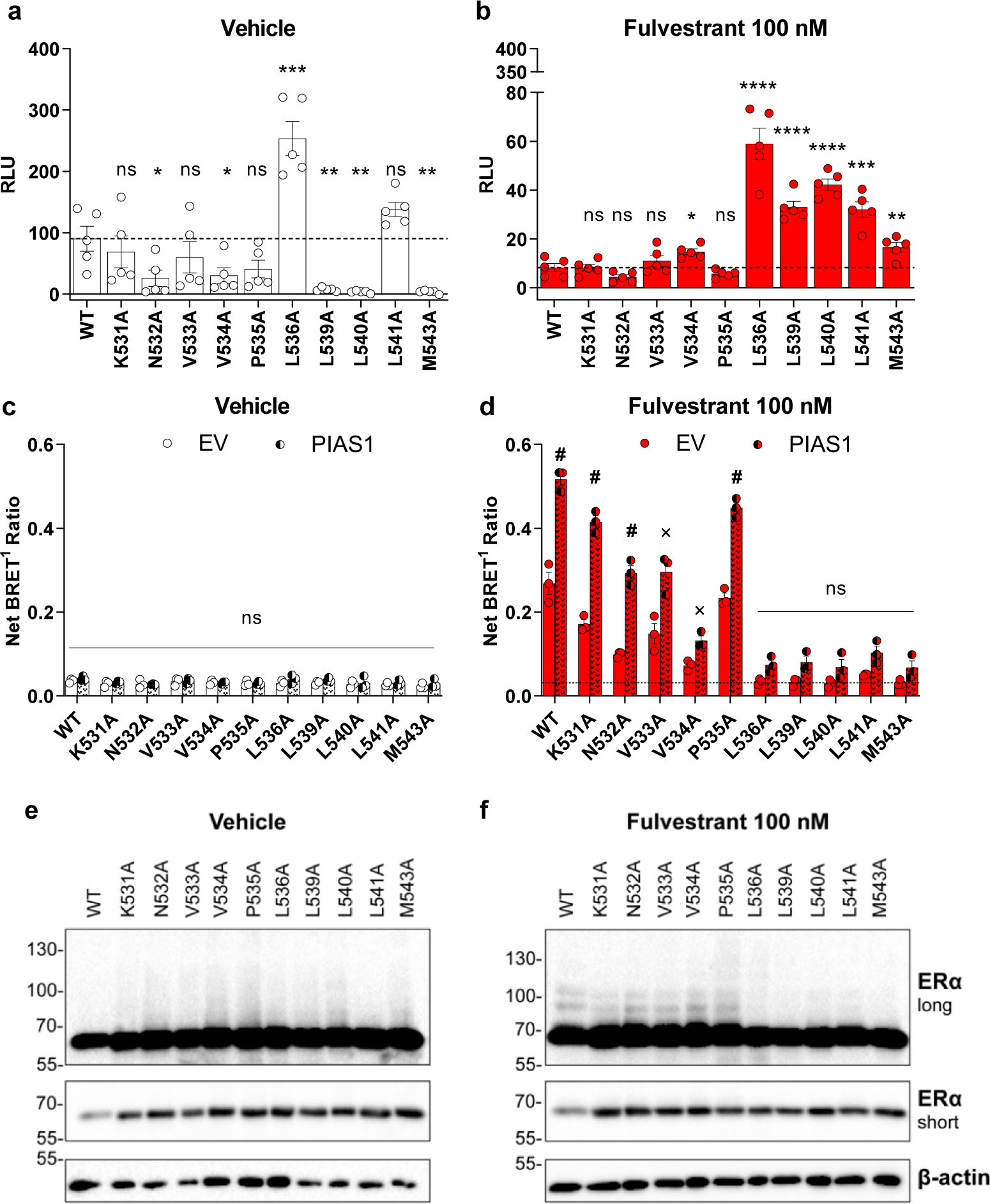
Importance of several residues located in helix 12 for fulvestrant-induced SUMOylation and transcriptional repression of ERα. (**a,b**) Transcriptional activity of ERα alanine mutants in the absence (a) or presence (b) of fulvestrant (Fulv) in an ERE3-TATA-LUC reporter assay in transiently transfected HEK293 cells treated with 100 nM Fulv for 24 h (N=3, mean values +/- SEM). Statistical significance in a Holm-Sidak multiple t-test is shown for the comparison between mutant at WT receptor (* P < 0.05, ** P < 0.01, *** P < 0.001; **** for p-value < 0.0001, “ns” P > 0.05). (**c,d**) Impact of alanine mutations in the absence/presence of transfected PIAS1 on ERα-SUMO3 interaction in transfected HEK293 cells treated with vehicle (c) or fulvestrant (d) for 2.5 h (N=3, mean values +/- SEM). Statistical significance in a Holm-Sidak multiple comparison test is shown for the comparison of BRET values in the presence vs absence of PIAS1 (# P < 0.05; x 0.05 < P < 0.1; “ns” p value > 0.1) (**e,f**) Impact of alanine mutations in H11-H12 loop or in the H12 helix on ERα modifications in HEK293 cells transfected as in a-b) and treated or not for 1h with 100 nM Fulv (N=3, representative blot shown).

Fulvestrant-induced transrepression was impaired with substitutions of H12 hydrophobic residues (L536, L539, L540, L541 and M543) in HEK293 cells (**Fig. 2b**), extending previous results ^20,25^. Point mutants in the H11-H12 loop maintained fulvestrant-induced transrepression, except for a slight induction in transactivation with V534A (**Fig. 2b**). In U2OS cells, increased activity was detected with mutations at positions 536, 539 and 540, while other mutations resulted in slighter gains in activity (**Supplementary Fig. 5b**). Thus, increased transcriptional activity in the presence of fulvestrant is observed even in mutants, such as L539A and L540A, which have reduced basal activity (**Fig. 2a,b**, **S5a,b**).

Next, we evaluated the impact of mutations on fulvestrant-induced coactivator recruitment and SUMOylation of ERα. In keeping with the lack of correlation between SERD activity and reduced coactivator recruitment, only minor gains in AF2-IM recruitment in the presence of fulvestrant were observed with some mutants with increased transcriptional activity (V534A, L536A, L539A, L541A and M534A), and none with L540A (**Supplementary Fig. 6**). Interestingly, several H12 mutants (L536A, L539A, L540A, L541A and M543A) that significantly reversed transrepression by fulvestrant (activity >2-fold that obtained with WT ERα) also had impaired fulvestrant-induced SUMO3 recruitment by BRET and Western analysis (**Fig. 2c,f**). The SUMO E3 ligase PIAS1 did not significantly boost fulvestrant-dependent BRET signals for these mutants, contrary to WT and other mutants, or to unstimulated cells (**Fig. 2c,d**). The V534A mutant, displays slight but significant gains in transcriptional activity in the presence of fulvestrant (**Fig. 2b**), also had low SUMO BRET signals, which were silghtly indiced by PIAS1 (**Fig. 2d**). Notably, the V534E mutation was previously reported to abolish SUMOylation and fulvestrant inverse agonism on estrogen target genes ^23^.

Together these results highlight the contribution of hydrophobic residues in H12 to ERα SUMOylation and transcriptional repression by pure antiestrogens.

### Hydrophobicity at position 536 is necessary for induced SUMOylation and maximal AE transrepression

To better understand the contribution of L536 to fulvestrant-induced SUMOylation, we characterized several mutants including L536R, P and Q, found in patients with metastatic ER+ breast cancer and associated with resistance to AE therapies ^39–45^. Substitutions for charged (R, K and D), polar (Q, S), short non-polar (G, A) residues, histidine (H) or proline (P) at position 536, led to increased basal activity of the mutant receptors with the ERE3-driven promoter (**Fig. 3a**). This observation is consistent with reported observations in HeLa cells for the L536G/A/K/P mutants ^38,46^. Hydrophobic residues Y, F, M, V at position 536 retained low basal activity (**Fig. 3a**). All mutations had residual activity compared to WT ERα in the presence of fulvestrant, with L536V, M and F being the least active (**Fig. 3b**). Fulvestrant-dependent SUMOylation was detected only with L536V/M/F in BRET assays in the presence or absence of PIAS1 (**Fig. 3 c,d**) and in Western analyses (**Fig. 3e, f**). Similar activity profiles were observed in ER-negative U2OS cells and SKBR3 breast cancer cells (**Supplementary Fig. 7**).

**Figure 3:**
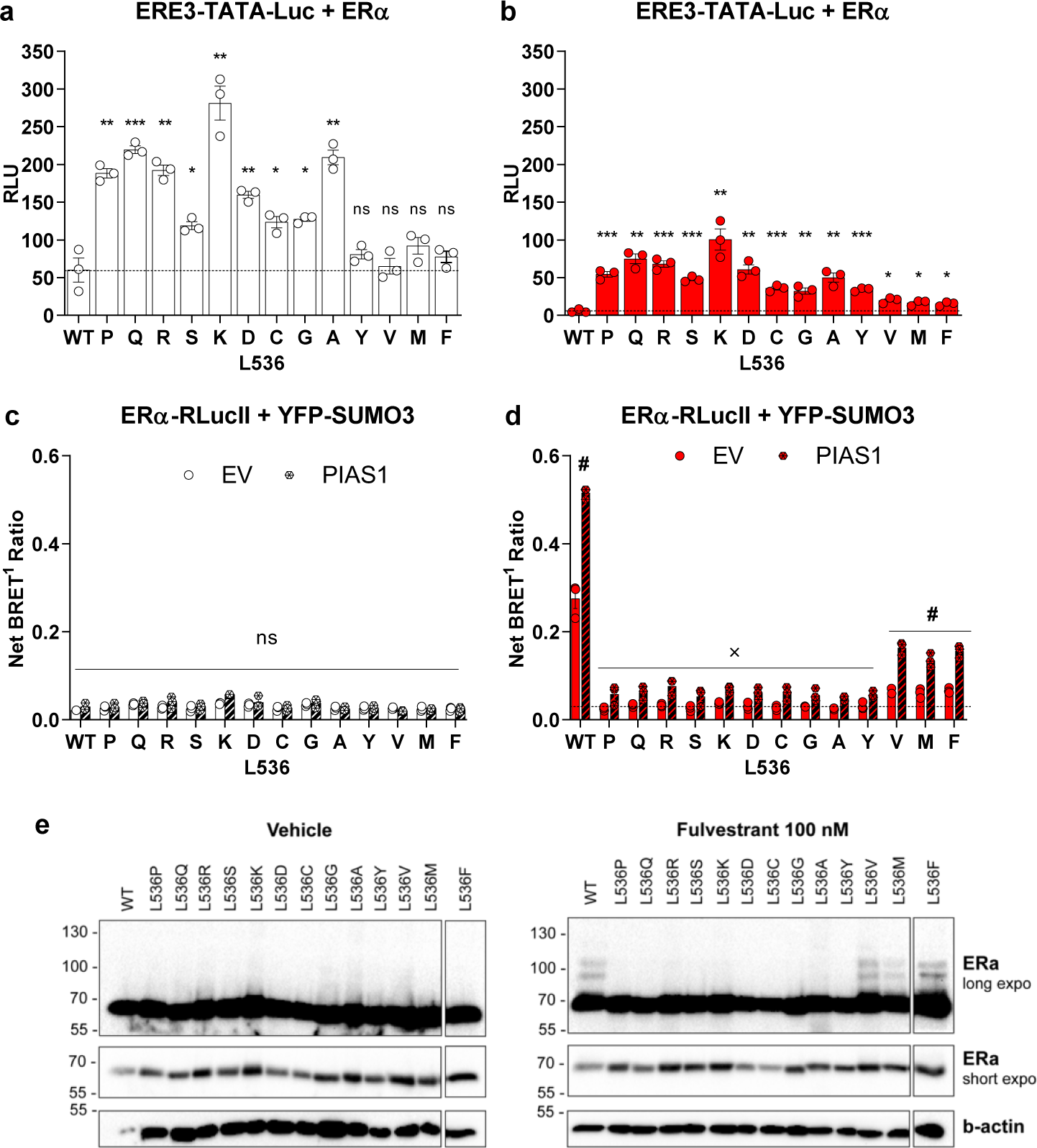
Hydrophobicity at position 536 is key for ERα SUMOylation and transcriptional suppression by fulvestrant. (**a,b**) Transcriptional activity of ERα L536 mutants expressed in transfected HEK293 cells in the absence (a) or presence (b) of Fulv (100 nM) (N=3; mean values +/- SEM; dashed line: RLU values for the WT receptor). Statistical significance in a in a Holm-Sidak multiple comparison test is shown for comparison between mutant vs WT receptor. * P < 0.05, ** P < 0.01, *** P < 0.001, **** P < 0,000, “ns” P > 0.05 (**c,d**) Impact of L536 mutations on the SUMO3 BRET signal in the absence (c) or presence (d) of Fulv (100 nM). Cells were co-transfected with an expression vector for PIAS1 or the empty parental vector (EV) as indicated (dashed line in D: net BRET1 values without treatment and PIAS1 overexpression). Statistical significance in a Holm-Sidak multiple comparison test is shown for comparison in presence vs absence of PIAS1. x 0.01<P<0.05; # P < 0.000001. (**e**) Impact of L536 mutations on ERα modifications in HEK293 cells transfected as above and treated for 1h with Fulv (100 nM). A representative blot is shown (N=3).

We further tested the clinically relevant L536R mutant, as well as L536S, used to obtain crystal structures with several SERDs ^27,28^, with our AE panel (**Fig. 4a**). All AEs behaved similarly to OHT in luciferase reporter assays with these mutants, contrary to their inverse agonist activity with the WT receptor (**Fig. 4a**). As with fulvestrant, induction of ERα SUMOylation was abolished by these mutations for all antiestrogens in BRET assays even with PIAS1 overexpression (**Fig. 4b**), or in Western analysis in transfected HEK293 cells (**Fig. 4c**). Thus, substitution of L536 for non-hydrophobic residues such as arginine or serine not only increases basal activity but also suppresses induction of SUMOylation by all tested antiestrogens, correlating with loss of their capacity to repress transcription more efficiently than tamoxifen.

**Figure 4:**
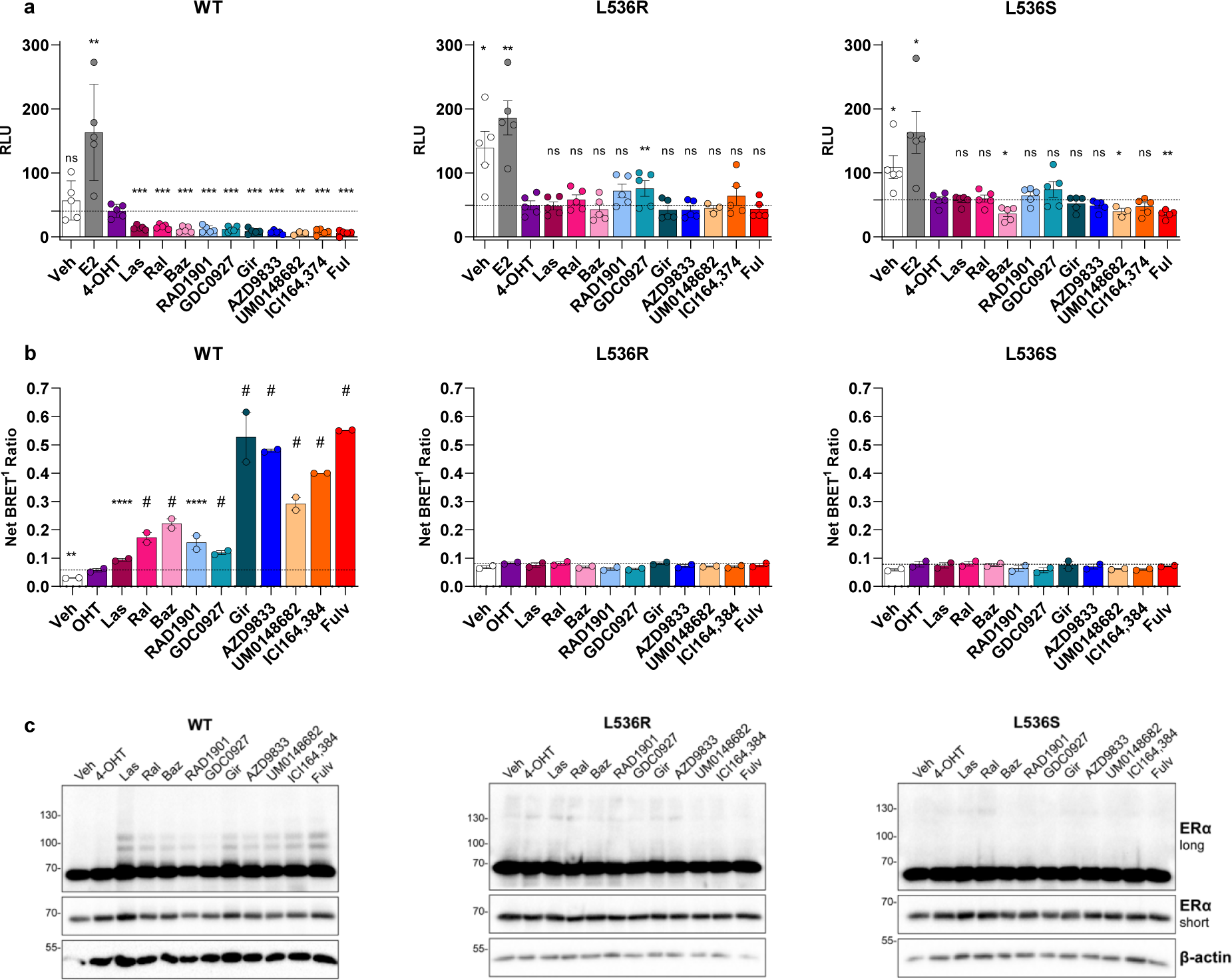
L536 is key for ERα SUMOylation and transcriptional suppression by different SERDs. (**a,b**) Impact of different classes of AEs on activity of L536R or L536S ERα mutants in luciferase reporter assays (a) or SUMO3 BRET assays with PIAS1 overexpression (b) in the presence of E2 (10 nM) or SERMs/SERDs (100 nM). (N=3; mean values +/-SEM; dashed lines: values obtained with the SERM 4-OHT). Statistical significance in a Holm-Sidak multiple comparison test is shown for the comparison between different AEs and 4-OHT. * P < 0.05, ** P < 0,01, *** P < 0,001, **** P < 0.0001; # P < 0.00001, “ns” P > 0,05. (**c**) Impact of L536R and L536S mutations on ERα modifications detected by Western analysis (N=3; representative blots shown).

### Pure antiestrogen side chains are unstable in L536S mutant ER LBD structures

So far, no crystal structures has been obtained of the WT ERα LBD complexed with SERDs. Stabilizing mutations including L536S have facilitated studying the impact of novel SERDs on the LBD structure ^27,28^. To understand the impact of long side chain AEs, we obtained cocrystals of ICI164,384 and analogs with variable side chain lengths with the ERα L536S LBD carrying stabilizing mutations C381S, C417S, C530S ^47^. Analogs included previously described C_10_(13), C_10_(14), C_10_(15), which have thioether linkers between the steroid core and side chains of length varying between 13 and 15 atoms ^25^ (**Supplementary Fig. 1**). Only C_10_(15) induced ERα SUMOylation and fully repressed transactivation ^25^, similar to desmethyl-ICI164,384 (UM0148682), the carbon analogue (**Supplementary Fig. 1**). Structure resolutions ranged between 1.5 and 2.2 Å. Asymmetric units (ASU) contain one monomer for ICI164,384 and desmethyl-ICI164,384, a dimer for C_10_(13), and two dimers for C_10_(14) and C_10_(15). Overall, each structure shows SERD steroidal cores identically positioned in the ligand binding pocket and H12 occupying the AF-2 cleft. However, AE side chains adopt divergent vectors near position 536 (**Fig. 5a**). Like in the rat ERβ-ICI164,384 (PDB: 1HJ1) and the human ERα-RU39411 (PDB: 6VIG) structures, the steroidal core is rotated by 180° compared to estradiol (**Supplementary Fig. 8**).

**Figure 5:**
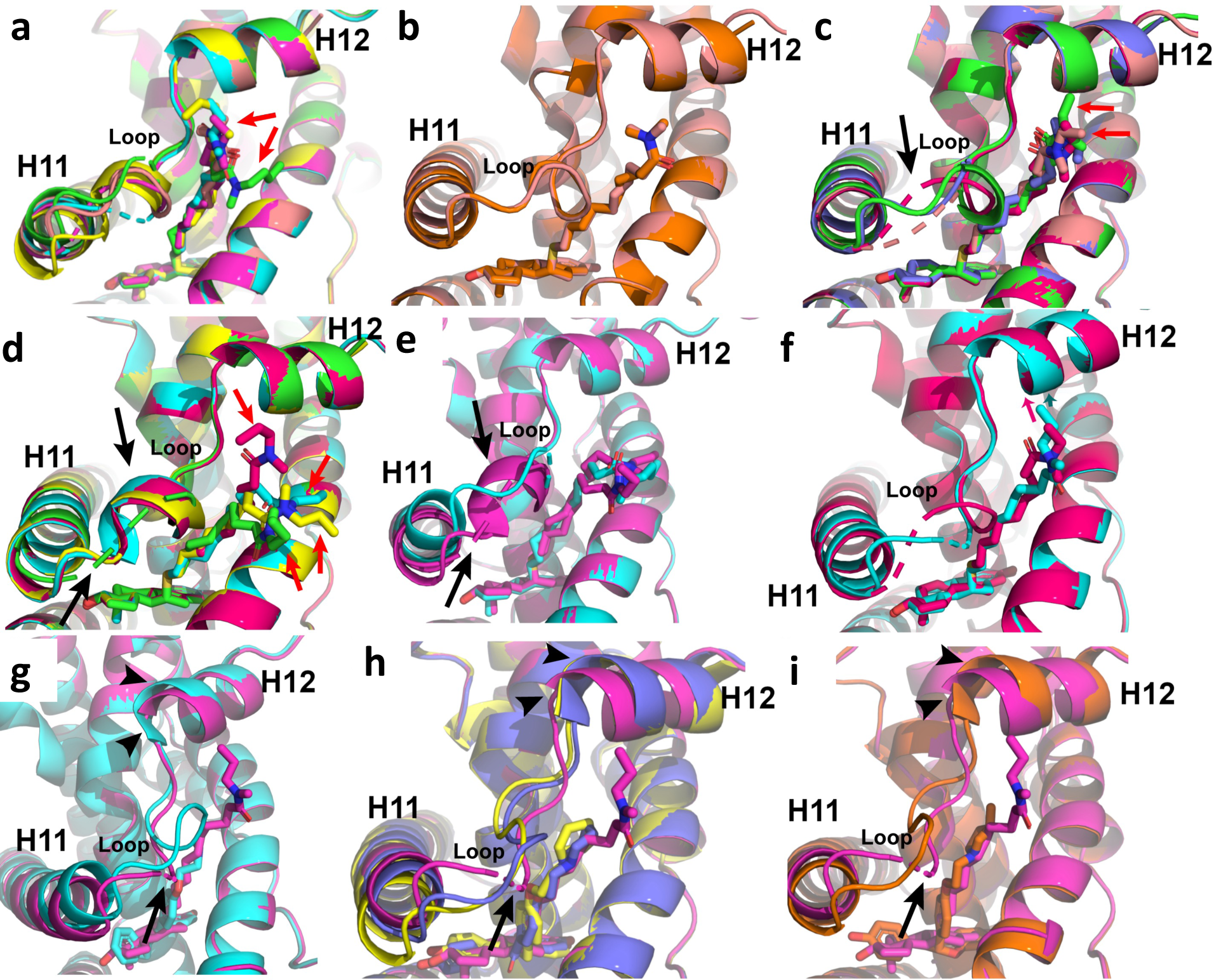
Aliphatic SERDs directly perturb ERα ligand binding domain helix 12. (**a**) Superposition based on alpha carbon positions of representative ERα monomers with different aliphatic SERD including C_10_(13) (beige), C_10_(14) (fuchsia), C_10_(15) (yellow), desmethyl ICI164,384 / UM0148682 (cyan), and ICI164,384 (green). (**b,c,d**) Superposition of each monomer in the asymmetric unit (ASU) of the ERα LBD in complex with C_10_(13) (**b**), C_10_(14) (**c**), C_10_(15) (**d**). (E) ERα LBD-ICI164,384 (cyan) superimposed with two representative monomers of the ERα LBD-C_10_(15) (fuchsia) structure. (**f**) ERα LBD-desmethyl ICI164,384 (cyan) superimposed on the ERα LBD-C_10_(14) (fuschia). (**g**) Superposition of ERα bound to endoxifen (cyan, PDB: 5W9D) and ERα-desmethyl ICI164,384 (fuchsia). (**h**) Superposition of ERα in complex with the SERM/SERDs Ral (blue, PDB: 7KBS), Baz (yellow, PDB: 4XI3), and desmethyl ICI164,384 (fuchsia). (**i**) Superposition of ERα LBD in complex with the oral SERD OP1074 (orange, PDB: 5UFX) and desmethyl ICI164,384 (fuchsia).

In the ICI164,384 thioether analogs, the relative atomic disorder within the structures correlates with increasing side chain length and with inverse agonism. The SERM C_10_(13) ^25^ is the best resolved structure, and the side chain with an N, N-dimethyl substituted terminal amide shows almost identical positioning near the H11-12 loop and H12 in both monomers (**Supplementary Fig. 9a**). C_10_(14), the N-ethyl-N-methyl analogue, shows poorer resolution than C_10_(13) (**Supplementary Fig. 9b**). Superposition of each monomer in the ASU confirms that C_10_(13) is more disordered as it gets closer to helix 12 (**Fig. 5c**, red arrows). N-methyl-N-propyl substituted C_10_(15) exhibits the least resolved electron density of the thioether series (**Supplementary Fig. 9c**). Superposition of each monomer in the ASU shows that C_10_(15) is highly disordered near the amide moiety (**Fig. 5d**, red arrows). Based on the positioning of the A monomer, C_10_(15) is of sufficient length to sample the AF-2 cleft. However, H12 binding in the AF2 cleft is likely more favorable for crystal formation, blocking ligand access. The ICI164,384 side chain, which adopts a similar orientation to those of molecules in C_10_(15) monomers B,C,D, is also poorly resolved as in most of the C_10_(15) monomers, suggesting a high degree of disorder (**Fig. 5e, Supplementary Fig. 8e**), while the C_10_(15) analog desmethyl-ICI164,384 reaches closer to H12 and is overall better resolved, except at the end of the side chain (**Fig. 5f, Supplementary Fig. 8d**).

The H11-12 loop is fully resolved in both monomers of the C_10_(13) structure, indicating relative stability (**Fig. 5b**), while C_10_(14) and C_10_(15) differ significantly in the H11-12 loop between the monomers in the ASU, with some adopting a pseudo-alpha helical character between positions 530 to 533 (**Fig. 5c,d**, black arrows). In the desmethyl-ICI164,384 structure, positions 531 and 532 in the H11-12 loop are poorly resolved (**Fig. 5f**) and in the ICI164,384 structure, position 532 is poorly resolved and the end of H11 is pushed up towards H12 (**Fig. 5e**). These data suggest that an ethyl substitution on the amide of the aliphatic side chain is sufficient to disorder the H11-12 loop.

H12 is also significantly altered in the structures of aliphatic SERDs compared to representative AEs within the SERM-SERD spectrum of activities. The SERM endoxifen shows a fully resolved H11-12 loop, H12 is slightly distant from the ligand binding pocket, and the helical character of H12 begins at position 536 versus 537 for the long-chain SERDs (**Fig. 5g**) ^48^. In the structures with bazedoxifene (Baz) and raloxifene (Ral), both associated with limited SUMOylation and ERα degradation, the H11-12 loop is also more ordered and H12 pushed slightly out of the AF-2 cleft (**Fig. 5h)** ^49,50^. The aliphatic SERDs also show an altered H12 and a greater variation in the H11-12 loop compared with the shorter side chain SERD OP1074 ^29^ (**Fig. 5i**). Together, these data suggest that aliphatic SERD side chains destabilize the H11-12 loop and influence H12 positioning.

### Molecular dynamics indicate length-dependent antiestrogen side chain mobility in ER*α* LBD structures with H12 in the coactivator binding groove

To explore potential perturbations associated with replacement of S536 by other residues in the presence of SUMOylation-inducing antiestrogens, we performed molecular dynamics simulations starting from the desmethyl-ICI164,384 co-crystal structure. We first docked fulvestrant into this structure and performed molecular dynamics simulations for both structures for 500 ns to ensure stability. The stabilized structures (frames extracted after 450 ns simulation time) were then used as a starting point to replace S536 with L (as in WT ERα) or R (a naturally occurring mutant) and subjected to another 500 ns of MD simulations. Backbone Root Mean Square Deviation (RMSD) analysis indicates that all systems were rapidly stable (<1.5 Å variation, **Supplementary Fig. 10**). Root Mean Square Fluctuations (RMSF) values for protein side chain residues in the H11-H12 loop and H12 helix (residues 529-546) fluctuated by less than 4 Å in all structures, indicating the absence of major delocalization of H12, irrespective of the mutations introduced or of replacement of desmethyl-ICI164,384 by fulvestrant. Side chain fluctuations were position-specific and influenced by the residue at position 536. Residues in the H11-H12 loop and at the C-terminus of H12 fluctuated most. At the H12 N-terminus, P535 and L540 side chains were relatively stable in all systems, with higher fluctuations of L536-539. Replacement of S536 by L or R increased fluctuations of H12 residues 536-539 with desmethyl-ICI164,384 and to a lesser degree with fulvestrant, and residues L541 and M543 also fluctuated more widely in the R536 model with desmethyl-ICI186,384; **Supplementary Fig. 11a)**.

RMSF values over the 500 ns simulation period for non-hydrogen atoms in each ligand indicated increasing fluctuation in the side chain with distance from the core in all systems, reaching 2 Å at position 25 (5 atoms from the core). The bulkier end of the side chain of fulvestrant was more mobile than that of desmethyl-ICI164,384. With both ligands, side chain mobility was increased in the R536 models. Of note, the fulvestrant side chain was more stable with the L536 model than with the S536 or R536 ones between side chain atoms 6-15 (positions 28-41, >1 Å difference), with 3-4 Å differences around the sulfinyl group (**Supplementary Fig. 11b**), suggesting stabilizing interactions.

### Amino acid identity at position 536 modulates interactions between the antiestrogen side chain and the N-terminus of H12 positioned in the coactivator binding groove

Mapping distances between the desmethyl-ICI64,384 amide methyl carbon (#34) in the S536 model and the CA atom of S536 and L539 in H12 and of L354 in H3 indicated that the ligand side chain terminus remains mostly within 5-10 Å of these residues throughout the simulation (**Fig. 6a, Supplementary movie 1**), the end of the side chain adopting a variety of conformations. In the L536 model, the desmethyl-ICI164,384 side chain extends close to L536 but more distal to L539 and L354 (**Fig. 6b, Supplementary movie 2)**. With the S536R replacement, the desmethyl-ICI164,384 side chain was relocated away from H12, and distances to all three residues increased to 15-20 Å (**Fig. 6c, Supplementary movie 3)**. Impressively, after 123 ns the R536 side chain relocated to form an ionic interaction with D351 on H3, closing the co-activator binding groove while the C-terminal part of H12 pivoted away from H3, explaining the increased RMSF values for H12 residues (**Supplementary Fig. 11d**). This was accompanied by interaction between the side chain carbonyl oxygen and the side chain hydroxyl group of Ser530 in the H11-H12 loop (interaction fraction via H-bond or water mediated contact >35%, **Supplementary Fig. 12c**).

**Figure 6:**
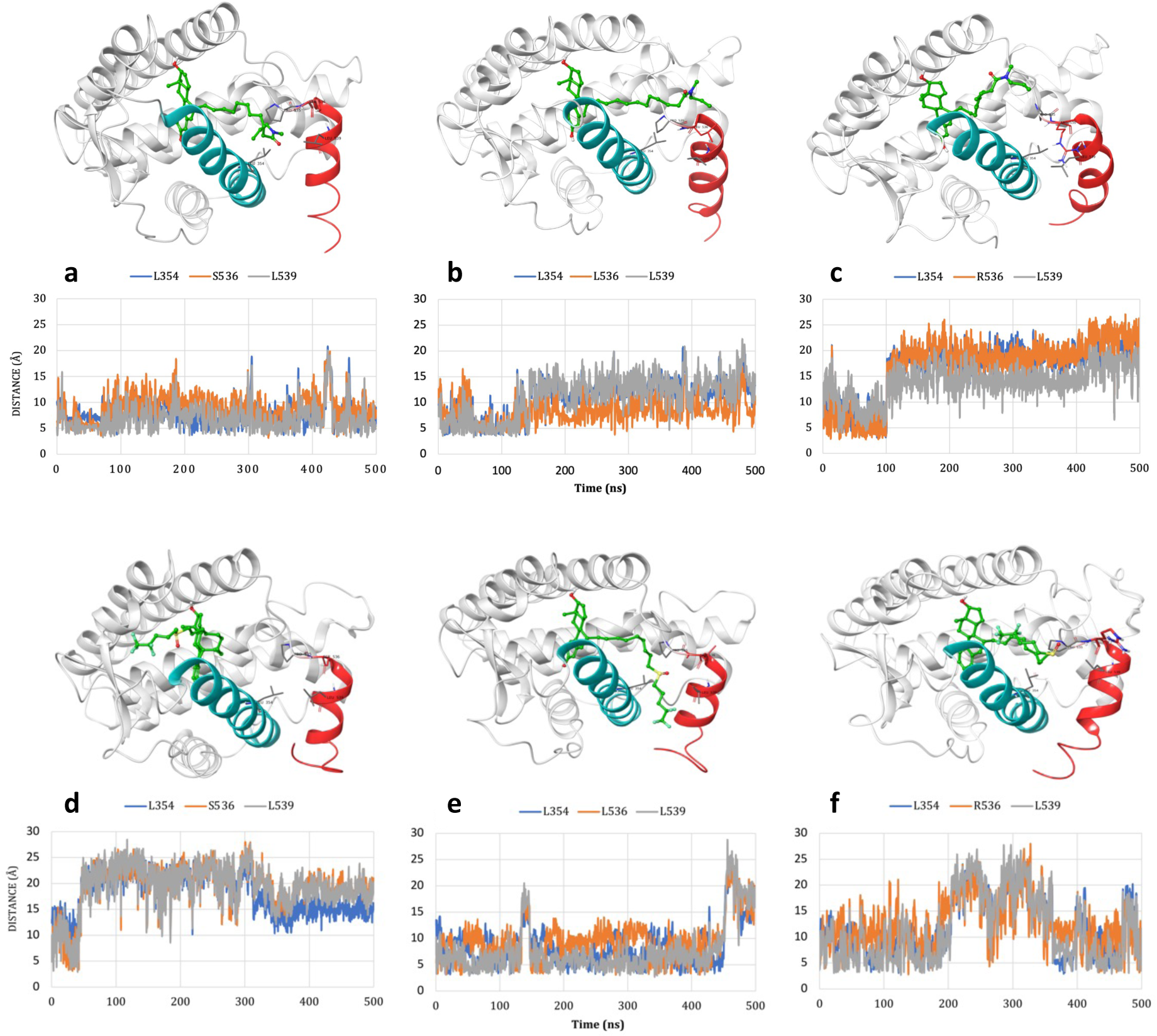
Mutations at position 536 affect proximity of the end of the ligand side chain to the N-terminus of H12. The plots provide the distance of ligand side chain atoms to the CA atom of residues L354, L539 and S536 (**a**, **d**), L536 (**b**, **e**) or R536 (**c**, **f**) during the 500 ns simulation. For desmethyl ICI164,384 (**a,b,c**), the distance is plotted from the amide methyl carbon (C34); for fulvestrant (**d,e,f**, the distance is plotted from the F36 atom.

With fulvestrant (**Fig. 6d-f)**, **Supplementary movie 4**), the side chain consistently projected towards H12 only in the L536 model, remaining close to L354 and L539 and extending into the coactivator binding groove (**Fig. 6e, Supplementary movie 5)**, resulting in the lower RMSF of the fulvestrant side chain in this system. The sulfinyl group recurrently interacted with the backbone amine of L536 (interaction fraction via H-bond 38% and water-mediated contact > 7%, **Supplementary Fig. 12**). The L536 side chain was mobile in solvent. On the other hand, in the S536 and R536 model, the end of the fulvestrant side chain fluctuated extensively (high RMFS, **Supplementary Fig. 11**) and often pointed away from H12 and towards the N-terminus of H3 and the C-terminus of H11 (**Fig. 6f, Supplementary movie 4&6)**.

Altogether, these results indicate that under crystallographic and molecular dynamics simulation conditions, H12 remains associated with the coactivator binding groove but that its positioning is variable. The antiestrogen side chain end is highly mobile and can associate transiently with the coactivator binding groove even with H12 positioned in the groove, in a manner favored by interactions between a hydrogen bond acceptor (sulfinyl or carbonyl) and the backbone amine at position 536, dependent on the nature of the amino acid at this position.

## DISCUSSION

Antiestrogens exhibit very different patterns of tissue-specific activity, which has been explained by different coactivator/co-repressor expression levels in the various estrogen-responsive tissues ^51^. The partial agonism of OHT is dependent on the AF1 activity of ERα, which recruits coactivators including the NCOA1/2/3 family ^32,35,52^. It has been postulated that pure antiestrogens can suppress AF1 as well as AF2 activity, but the mechanisms of this suppression remain uncharacterized. In this respect, our observations that, similar to 4-OHT, most AEs induce recruitment of the AF1-interacting domain of the SRC1 coactivator by full-length ERα in BRET assays in live cells, were unexpected. While we cannot exclude that different AF1-binding coactivators may be less efficiently recruited by pure AEs, this finding suggests that other mechanisms may underly the full antagonist activity of fulvestrant.

Reported impacts of fulvestrant include induction of ERα insolubility, intra-nuclear immobilization, modification by SUMOylation, reduced interaction with DNA as well as ubiquitination and degradation ^15,21–23^. Some of these properties may be linked, since SUMOylation may result in changes in protein-protein interactions affecting mobility and extractability, and can trigger ubiquitination by SUMO-targeted E3 ubiquitin ligases ^53^, resulting in increased proteasomal degradation. In addition, SUMOylation at EREs preceded rapid loss of ERα DNA binding in genome-wide ChIPseq studies and chromatin compaction ^23^. Results presented here indicate that transcriptional suppression by AEs correlates well with induction of SUMOylation, consistent with our previous observation of complete suppression of ERα transcriptional activity by fulvestrant in HepG2 cells in the absence of ERα degradation ^20^. On the other hand, correlation with ERα degradation in ER+ breast cancer cells was imperfect. The ARV-471 PROTAC, which induces efficient ERα degradation, did not increase SUMOylation and yielded transcriptional activity more comparable to that of 4-OHT than fulvestrant. SERDs with acidic side chains ^28,54^ also did not detectably induce SUMOylation in HEK293 cells (although partial induction was observed in MCF7 cells, not shown) and were less efficient than fulvestrant at suppressing residual transcriptional activity in these cells. Of note, AZD9496 and GDC0810 were observed to have uterotrophic activity in ovariectomized mouse models ^24^.

Notably, ERα SUMOylation monitored by BRET ^25^, complementing detection of receptor modification in Western analyses, revealed a spectrum of efficacies of ERα SUMOylation by AEs. ICI164,384 and oral SERDs Giredestrant (Gir) and Camizestrant (AZD9833), currently in phase III trials, were the most efficacious at inducing SUMOylation. Gir and AZD9833 have different scaffolds but share a side chain longer than that of structurally-related AEs with lower SUMOylation efficacies, consistent with the reported importance of the AE side chain length in ER antagonism ^19,29^ and induction of ERα modifications ^25^.

Point mutations in the H11-H12 loop and on H12 can impact ERα transcriptional activity in response to AEs ^20,22,25,29,38,40,45,49^. Our alanine scanning mutagenesis further indicated that L536, L539 and L540 in H12 contribute to fulvestrant-induced ERα SUMOylation as well as transcriptional repression. While Ala substitutions in the H11-H12 loop had a limited impact on SUMOylation, replacing V534 by non-hydrophobic residues also abrogated SUMOylation and efficient transcriptional repression ^23^. Together, these studies indicate that hydrophobicity at several positions in H12 or close to the H12 N-terminus contributes to complete antagonist activity.

The H11-H12 loop may act as a spring keeping H12 away from its active conformation due to the aqueous exposure of hydrophobic residues V533, V534, P535 and L536 ^45,55^. Indeed, replacement of L536 by charged (R, H), polar (Q) or less hydrophobic residues (P), expected to reduce this spring-like action, increased basal activity ^45,55^. While the constitutive activity of more polar L536 mutants could reduce affinity for pure antiestrogens, repression of basal constitutive activity comparable to that of OHT at 100 nM in reporter assays supports maintained binding capacity and lack of SUMOylation was observed even at elevated doses of fulvestrant (1-5 μM). Further, the L536Y, L539A and L540A mutants, which did not display constitutive activity in our assays, also exhibited minimal SUMOylation and a significant increase in activity with fulvestrant.

As we did not obtain crystal structures of WT ERα LBD with fulvestrant or ICI164,384, we used, as previously done for other SERDs ^27,28^, the S536 LBD mutant also carrying C381S, C417S and C530S substitutions, which favor the antagonist conformation of ERα even in the presence of agonists ^47^. Mutation S536 may stabilize the H12 N-terminus structure in presence of pure antiestrogens in part through an observed H-bond formed with the backbone amine of L539. In our structures with the ERα S536 LBD, H12 occupies the coactivator binding groove, possibly due to the stabilizing mutations. Structural analysis revealed that the side chains of aliphatic SERDs in complex with ERα S536 become disordered when their length exceeds 13 atoms, with corresponding disorder of the H11-H12 loop. Accordingly, ICI164,384 derivatives with side chains longer than 14 atoms were associated with induction of ERα SUMOylation and gain of pure antiestrogenicity (e.g., C_10_(15) ^25^ and desmethyl-ICI164,384) with wt ERα, suggesting that interference with H12 positioning in the coactivator binding groove is linked to recruitment of the SUMOylation machinery.

Consistent with observations from crystal structures, MD simulations indicated increasing fluctuations of the side chain atoms in a length-dependent manner with all models. The side chain sampled multiple conformations and interacted transiently with LBD surface residues including with H12 in the coactivator binding groove, in a manner affected by the residue at position 536. The fulvestrant side chain, and occasionally that of desmethyl-ICI164,384, extended into the coactivator binding groove of the L536 ERα LBD. Differing from the rat ERβ/ICI164,384 complex in which H12 is undefined ^56^, the sulfinyl group of fulvestrant interacted with the backbone amine of L536 in H12 positioned in the coactivator binding groove. The carbonyl group of desmethyl-ICI164,384 also engaged in hydrogen bonding with the same group, albeit less stably. Interestingly, the carbonyl group of short acidic side chain AE GW5638 engaged in a similar interaction in the wt ERα LBD and relocalized the N-terminus of H12 in a manner that resulted in increased exposure of hydrophobic amino acids ^54^. This interaction was not observed in the complex between another AE with acidic side chain, AZD9496, and the L536S LBD ^28^. Remarkably, both the stabilizing S536 and the unstable R536 mutations, which respectively replace the hydrophobic Leu side chain by a polar or charged residue, resulted in relocation of ligand side chains away from H12 during most of the simulation time (with fulvestrant for S536 and with both ligands for R536). In the R536/desmethyl-ICI164,384 complex, establishment of a salt-bridge between the basic side chain of R536 and D351 on H3 resulted in closure of the coactivator binding groove. Together, our results suggest that positioning of the AE side chain towards the coactivator binding groove and interaction with the N-terminus of H12 occupying the groove correlate with induced SUMOylation and optimal antiestrogenicity. We speculate that the increased local exposure of hydrophobic determinants (H12 long hydrophobic amino acids and possibly also the end of the AE side chain positioned in an adjacent manner) is important for recognition by the SUMOylation machinery, such as E3 SUMO ligases. Whether these interactions ultimately lead to displacement of H12 from the coactivator binding groove remains impossible to assess at this point.

While fulvestrant was the most efficacious antiestrogen in inducing SUMOylation in cell-based assays, its poor oral bioavailability is thought to diminish its clinical efficacy. We find that more bioavailable SERDs under current Phase II/III trials induce SUMOylation to varying degrees, in addition to their capacity to induce ERα insolubility and intra-nuclear immobilization ^22^. Whether the negative impact of L536 mutations on SUMOylation reduces the clinical efficacy of SUMO-inducing SERDs in tumors carrying these mutations remains to be determined in models assessing acquired long-term response to these agents. Nevertheless, our results demonstrate that induction of SUMOylation, potentially paralleled by increased ubiquitination, contributes to the efficient antagonistic profiles of fulvestrant and new generation SERDs, and indicate that the capacity of long aliphatic AE side chains to extend to the coactivator binding groove close to the N-terminus of H12 in its antagonist position is a key component of complete antiestrogenicity.

## Methods

### Cell lines and reagents

Cell lines were obtained from the American Type Culture Collection (ATCC) and maintained in a humidified 37°C, 5% CO_2_ incubator. MCF-7 cells were maintained in alpha modification of Eagle’s medium (αMEM, Wisent) supplemented with 10% Fetal Bovine Serum (FBS, Sigma) and 2 mM L-glutamine (Wisent). HEK293T, U2OS and SKBR3 cells were cultured in Dulbecco’s modified Eagle’s media (DMEM, Wisent) supplemented with 10% FBS. Forty-eight hours before each experiment, cells were switched to DMEM without phenol red, supplemented with 10% charcoal-dextran-striped FBS (Sigma) and 4 mM L-glutamine.

The transfection reagent polyethylenimine (PEI) was ordered from Polysciences, Inc. 17β-estradiol (E2) and lasofoxifene (Las) were purchased from Sigma, 4-hydroxytamoxifen (4-OHT), raloxifene (Ral) and bazedoxifene (Baz) were obtained from Tocris, fulvestrant (Fulv), GDC0927, GDC9545 (Gir), AZD-9833 and RAD-1901 were purchased from MedChemExpress. ICI164,384 and its derivative desmethy-ICI164,384 were synthesized at the IRIC medicinal platform (Supplementary Material).

Rabbit monoclonal anti-ERα, clone 60C (04-820) and mouse monoclonal anti-β-actin, clone AC-15 (A5441) were purchased from EMD Millipore and Sigma respectively. Horseradish peroxidase (HRP)-conjugated secondary antibodies were obtained from Jackson. Polyvinylidene difluoride (PVDF) membranes were purchased from EMD Millipore. Enhanced chemiluminescence (ECL) detection reagents were ordered from Biorad.

ERα mutants in BRET donor vector pcDNA3.1-ERα-RLucII (generated as described in ^35^) were generated by site-directed mutagenesis (Table S1). ERα mutants were generated by site-directed mutagenesis using pSG5-HEG0 expressing vector coding for the WT human receptor (gift from P. Chambon, IGBMC, France). Acceptor vectors L2N2-YFP, which contains two LXXLL motifs fused to the Venus YFP, and AF1ID-YFP, which contains the AF1 receptor interacting region of coactivator NCOA1/SRC1 (glumatine-rich region) fused to the YFP Topaz, were described before ^35^. Acceptor pEYFPC1-SUMO3 was obtained from M. Dasso (National Institutes of Health, Bethesda, MD) ^57^. The reporter vector for luciferase reporter assays was obtained by cloning three copies of the estrogen response element consensus sequence (AGGTCA CAG TGACCTA) in the pGL3 vector containing the firefly luciferase reporter gene. pGFP-N2 expression vector was obtained from Clontech.

### Bioluminescence Resonance Energy Transfer (BRET) Assays

HEK293T cells were switched to estrogen depleted media 48h prior to each BRET experiment. For assays in the absence of PIAS1 overexpression, cells were co-transfected with 15 ng of pcDNA3.1-ERα-RLucII (donor) either alone (for background evaluation) or together with 900 ng of coactivator vectors or 600 ng of SUMO3 vector (acceptors) in 75 μl of PBS and completed to 1.5 μg with salmon sperm DNA. Transient transfections were performed using PEI (3 µg of linear PEI for each µg of DNA diluted in PBS) that was mixed with DNA (1V/V, 150 ul total) and left for 12 min at room temperature.

For BRET assays in the presence of PIAS1, cells were co-transfected with 50 ng of pcDNA3.1-ERα-RLucII (donor) either alone (for background evaluation) or together with 500 ng of SUMO3 vector (acceptor) and 200 ng of pCMV-Flag empty vector (EV) or an expression vector for PIAS1 in 50 μl of PBS and completed to 1 μg with salmon sperm DNA. Transient transfections were performed using PEI (3 µg of linear PEI for each µg of DNA diluted in PBS) that was mixed with DNA (1V/V, 100 ul total) and left for 12 min at room temperature.

Cells in suspension (1.25 x 10^6^ cells / ml) were added to the PEI:DNA solution and 100 μl of this mixture was seeded in 96-well white-bottom culture plates (Corning). Two days after transfection, medium was replaced by Hank’s Balanced Salt Solution (HBSS) supplemented with 4,5 g/L dextrose and cells were treated with E2 (10 nM) or tested compounds (100 nM) in triplicates for 45 min (coactivator BRET) or 2h30 (SUMO3 BRET) before BRET assays. Coelenterazine-H (Nanolight Technology) resuspended at 2 mM in 100% ethanol was added to each well to a final concentration of 5 µM and incubated for 5 min at room temperature protected from light. Readings were then collected using a MITHRAS LB940 (Berthold Technology) multidetector plate reader, allowing the sequential integration of the signals detected in the 485/20 nm and 530/25 nm windows, for luciferase and YFP light emissions, respectively. The BRET signal was determined by calculating the ratio of the light intensity emitted by the YFP fusion over the light intensity emitted by the Luc fusion. Net BRET values represent the difference between BRET ratios obtained with ERα fused to RLucII and values obtained with unfused RLucII.

### Luciferase reporter assays

HEK293T, U2OS and SKBR3 cells were switched to estrogen-depleted media 2 days before each experiment. Transient transfection of HEK293T and U2OS cells were performed using polyethylenimine (3 μg linear PEI per μg DNA in PBS). Cells in suspension (7.5 x 10^5^ cells in 1.35 ml PBS 1X) were co-transfected with the luciferase reporter vector pGL3-ERE3-TATA-Luc (1.25 μg), the pSG5-ERα expression vector or parental empty vector (0.6 μg), and the pGFP-N2 internal control expression plasmid (100 ng).

SKBR3 cells were transiently transfected by electroporation. Cells were resuspended at 20 x 10^6^ cells per ml in electroporation media (complete culture media supplemented with 15 mM HEPES pH 7.0). For each condition, 5 x 10^6^ cells were added to a DNA mix containing the luciferase reporter vector pGL3-ERE3-TATA-Luc (2 μg), the pSG5-ERα expression vector or empty parental vector (1 μg), a GFP internal control expression plasmid (0.5 μg) and salmon sperm DNA to 40 ug DNA total. This mixture was transferred into an electroporation cuvette (Biorad) and electroporated at 240V/950 μF using a Gene Pulser Transfection Apparatus (Biorad). Cells were pelleted and resuspended at 1M cells/ml in culture media and aliquoted in 96-well plate (1 x 10^5^ cells per well).

Twenty-four hours after transfection, cells were treated for another 24h with either 10 nM E2 or 100 nM antiestrogens or vehicle (DMSO). GFP signals were measured prior to luminescence assessment with a Flexstation microplate reader (Molecular Devices) at an excitation/emission of 400/505 nm. Cells were assayed for luciferase activity in the presence of a luciferin lysis solution [30 mM trycine, 1,61 mM (MgCO_3_)_4_•Mg(OH)_2_•5H_2_O, 4 mM MgSO_4_, 0.15 mM EDTA pH 7.0, 50 mM DTT, 0.81 mM coenzyme A, 0.7 mM D-luciferin, 0.8 mM ATP, 1% Brij58, 100 mM Tris, 20 mM (CH_3_COO)_2_Mg and 10 mM EGTA in 1X PBS]. Luminescence measurements were acquired after 10 min incubation at room temperature using a MITHRAS LB940 multidetector plate reader (Berthold Technology). Results were normalized to GFP expression. The relative luciferase signal (RLU) was obtained by normalizing the luciferase signal to that of the control condition (transfected only with the reporter vector and GFP internal control).

### Western blotting

MCF-7 cells were switched to estrogen-depleted media and seeded in 6-well culture plates (5×10^5^ cells per well). HEK293 or U2OS cells that were transfected for luciferase assays were transferred to 12-well culture plates (2.3×10^5^ cells per well). Two days after, cells were treated for 1 h with 100 nM antiestrogens and harvested in ice cold 1X PBS. Cell pellets were resuspended in high detergent buffer (50 mM Tris-HCl pH 7.5, 150 mM NaCl, 5 mM EDTA, 2% SDS, 0.5% Triton X-100, 1% NP-40) supplemented with protease inhibitors (aprotinin, leupeptin, pepstatin, phenylmethylsulfonyl fluoride and N-Ethylmaleimide) for protein extraction. Samples were sonicated for 15 min, with 30 sec ON and 30 sec OFF cycles at high power in a water-bath sonicator (Bioruptor, Diagenodde). Proteins were denatured in 1X Laemmli buffer at 95°C for 5 min. Samples were loaded on an SDS-polyacrylamide gel (7% acrylamide, 20-50 µg protein/lane) and transferred to a PVDF membrane, then blotted overnight at 4°C with primary antibodies targeting either ERα or β-actin. After incubation with HRP-conjugated secondary antibodies, membranes were revealed with the ECL Clarity solution (BioRad). Signals were detected using a ChemiDoc gel imaging system (BioRad) or by film exposure (Denville Scientific Inc).

### Crystallography

The ERα LBD with C381S, C417S, C530S, and L536S mutations was expressed in *E.coli* BL21(DE3) and purified as described ^29^. Co-crystals were generated by incubating 10 mg/mL purified ERα LBD with 1 mM aliphatic SERD at 4°C overnight. Hanging-drop vapor diffusion was used for each structure using Hampton VDX plates (Hampton Research, HR3-140). All crystals were grown in 10-25% PEG 8,000, pH 6.5-7.5, and 200 mM MgCl_2_. Clear hexagonal crystals emerged at room temperature between 24 and 72 hours. 25% PEG 8,000 was used as cryo-protectant for all crystals. All X-ray data sets were collected at the Advanced Photon Source, Argonne National Labs, Argonne, Illinois on the SBC 19-BM beamline (0.97 Å). **Table S2** contains data collection and refinement statistics. **Supplementary Fig. 8** shows representative 2mFo-DFc difference maps for the aliphatic SERDs in the ligand binding pocket.

### Molecular dynamics simulations

The crystal structure of the desmethyl-ICI164,384 with estrogen alpha was preprocessed using the protein preparation wizard of Schrodinger (http://www.schrodinger.com/). The processed structure was subjected to energy minimization using the OPLS4 force field in the impact module. The grid box for docking was created by selecting desmethyl-ICI164,384 using Glide. The ligand structure of fulvestrant was prepared with the Ligprep module of Schrödinger and for docking Glide XP docking method ^58^ was employed. Mutant systems were prepared using the residue scanning calculation workflow in Maestro. When residue serine 536 was mutated to leucine and arginine for both the ligands system respectively, structures were minimized using the prime module of Schrodinger.

Molecular dynamics (MD) simulations were performed on the primed structure of desmethyl-ICI164,384 and fulvestrant with S536, and the mutated S536L and S536R structures. Solvation of the model was done with the TIP4P model in a cubic periodic boundary box. The ensuing systems of approx. 34000 to 35000 atoms were minimized and pre-equilibrated by standard procedures implemented in Desmond ^59,60^. Periodic wall distance was set to greater than 10 Å, thereby avoiding direct interactions with the periodic images. Gradient threshold was set to 25 kcal/mol/Å using steepest descents energy minimization, followed by a convergence threshold of 1 kcal/mol/Å by the L-BFGS (Low-memory BroydenFletcher-Goldfarb-Shanno quasi-Newtonian minimizer). The MD run was performed on the equilibrated systems at a constant temperature of 310 K and constant pressure for a duration of 500ns. The equations of motion were integrated with a 2 fs time step in the NPT ensemble. The SHAKE algorithm was applied to all hydrogen atoms, the van der Waals cut-off was 9 Å and the temperature was maintained at 310 K, employing a Nose–Hoover thermostat with a relaxation time of 1 ps.

### Statistical analyses

All experiments were performed at least 3 times independently. Statistical analyses were performed in GraphPad Prism using the Holm-Sidak multiple comparison test.

### Accession codes

Structures were deposited in the Protein Data Bank with the following codes:

## Acknowledgments

Funding for this study was provided by the Canadian Institutes for Health Research (grant # PJT156199) to SM and the Susan G. Komen Foundation (CCR19608597) to SWF. Results shown in this report are derived from work performed at the Structural Biology Center funded by the U.S. Department of Energy, Office of Biological and Environmental Research and operated for the DOE Office of Science at the Advanced Photon Source by Argonne National Laboratory under Contract No. DE-AC02-06CH11357.

## Contributions

MD performed most experiments and analyzed results, prepared figures and cowrote the manuscript. FS performed molecular dynamics simulations, interpreted results and cowrote the MD sections of the manuscript with DAS and AM. MEE contributed to experiments on the impact of L536 mutations on ERα SUMOylation and to analysis of results. KT performed reporter assays in U2OS cells and analyzed results. SW performed Fig. 1f, 2c,d and 3c,d. JP synthesized desmethyl-ICI164,384 under guidance from RMS and AM and prepared Figure S1. JG helped with interpretation of results. DAS supervised the design and interpretation of molecular dynamics. GG supervised the crystallography data acquisition and interpretation. AM supervised the chemistry and molecular dynamics work. SWF performed the crystallographic work and analysis and co-wrote the manuscript. SM conceived the study, supervised experiments, interpreted results and co-wrote the manuscript.

## EXTENDED DATA

### FILMS

**Desmethyl ICI164,384 with ERα S536 LBD**

**Desmethyl ICI164,384 with ERα L536 LBD**

**Desmethyl ICI164,384 with ERα R536 LBD**

**Fulvestrant with ERα S536 LBD**

**Fulvestrant with ERα L536 LBD**

**Fulvestrant with ERα R536 LBD**

**METHODS:** Synthesis of Desmethyl ICI164,384

## SUPPLEMENTARY MATERIAL

**SUPPLEMENTARY FIGS: Supplementary Figures S1-12**

**SUPPLEMENTARY TABLES: Supplementary Tables S1-S2**

