## Supplementary material for "Structural determinants of pure antiestrogenicity"

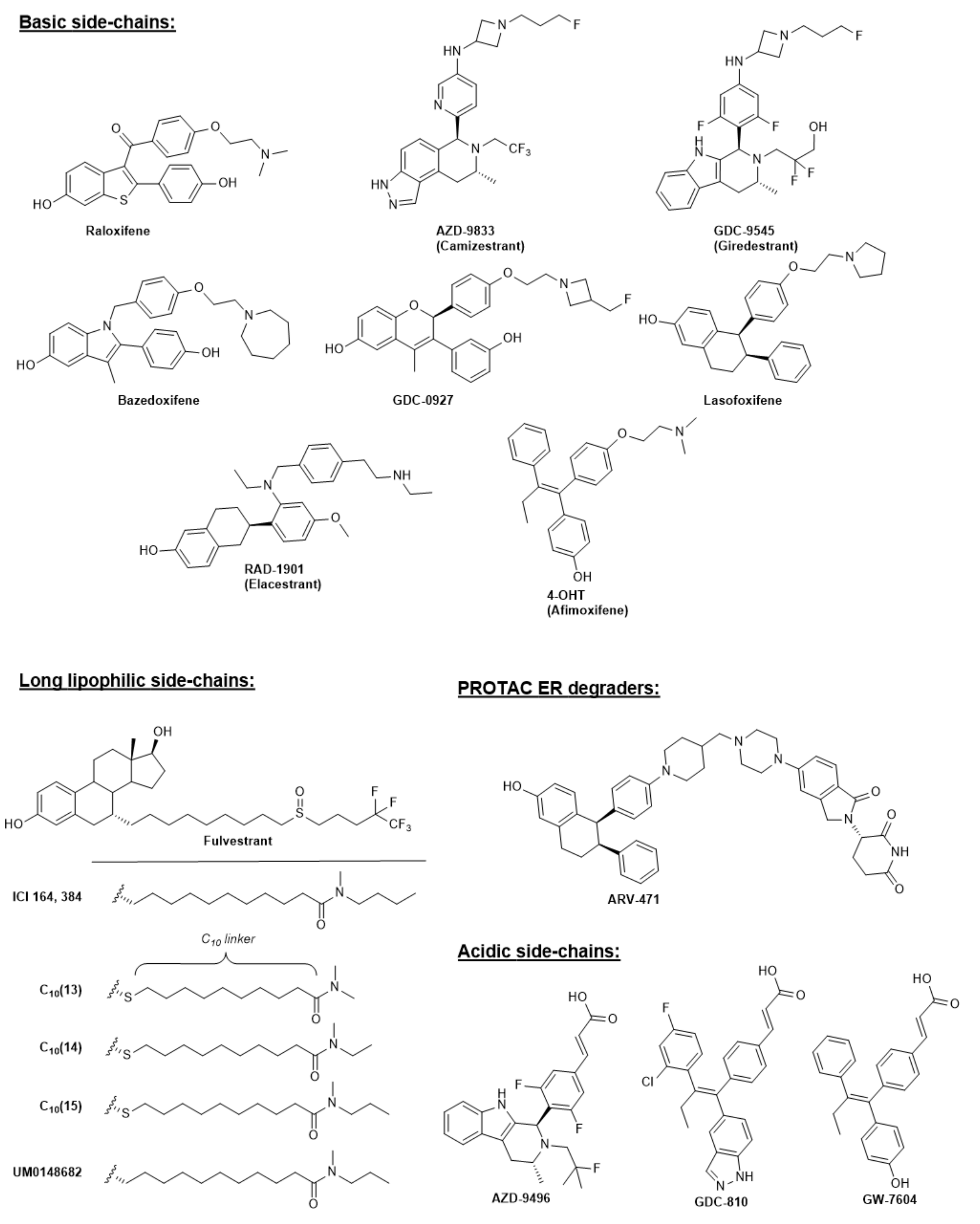


**Supplementary Fig. 1: Antiestrogens used in this study.** Antiestrogens are grouped by side chain type, either basic, long and mostly lipophilic, acidic, or interacting with the Ubi E3 ligase cereblon in the case of the ARV471 PROTAC.

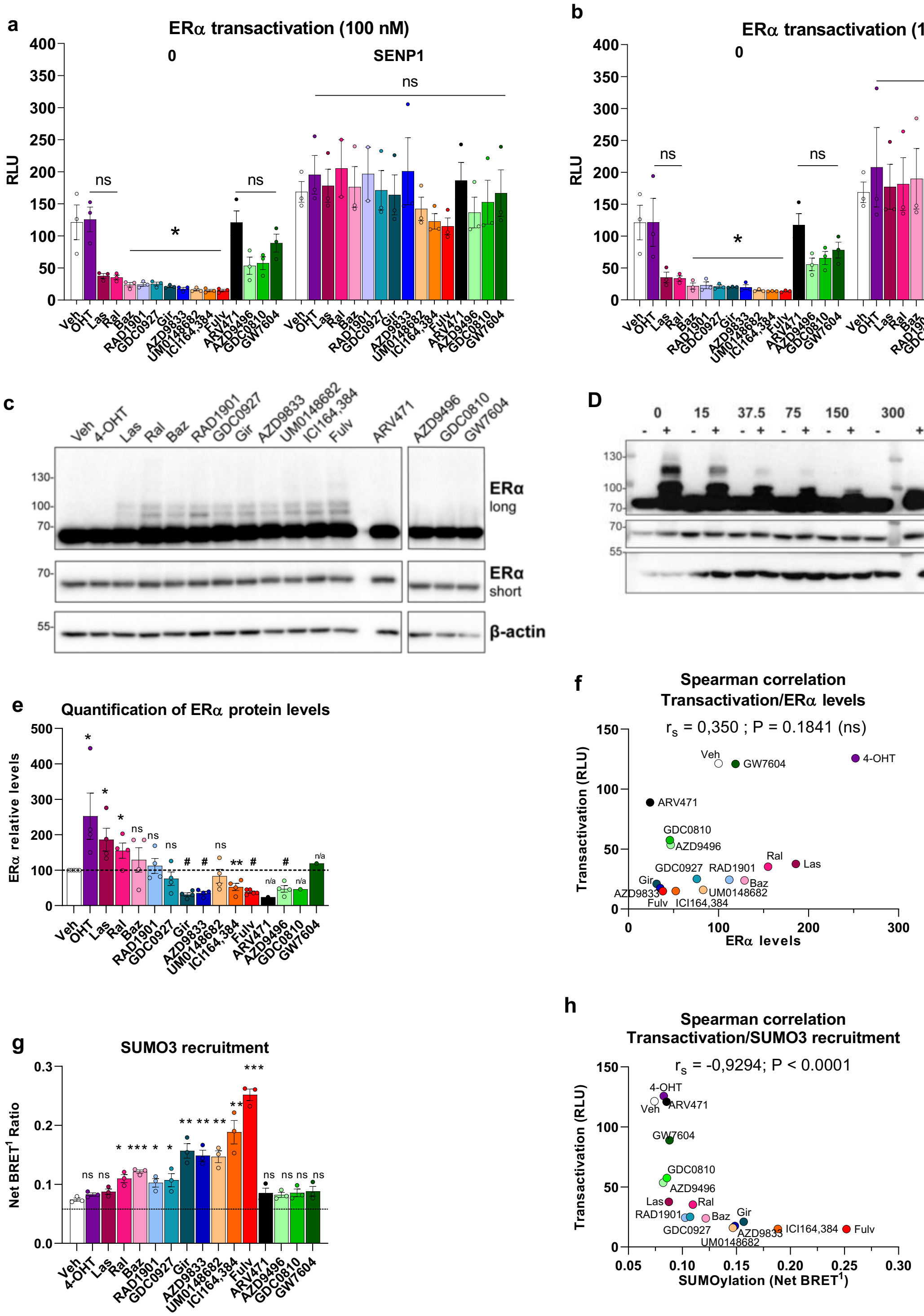


**Supplementary Fig. 2: The differential impact of AEs on ERα transactivation correlates better with SUMOylation than with degradation. A-B)** Differential transcriptional repression by SERMs and SERDs in HEK293 cells transiently transfected with the ERE3-TATA-LUC reporter vector together with an ERα expression vector supplemented or not with a SENP1 expression vector, and an expression vector for GFP for internal standardization. Cells maintained in estrogen-depleted medium were treated with estradiol (10 nM) or antiestrogens (100 nM (A) or 1 mM (B)) for 24 h. Values are means of 3N except for Ral and Baz with SENP1 (N=2), error bars represent the s.e.m. (* $p \leq 0.05$) **C)** Impact of antiestrogens on ERα modifications in a Western assay. HEK293 cells maintained for 2 days in estrogen-deprived media were transiently transfected with an ERα expression vector and were treated 48 h later with various antiestrogens for 1 h. (N=3, representative blot shown) **D)** Overexpression of the deSUMOylase SENP1 abolishes fulvestrant-induced modifications of ERα. HEK293 cells transiently transfected with an ERα expression vector together with variable amounts of a SENP expression vector (0-300 ng) complemented with salmon sperm DNA were treated with Fulv (100 nM) for 1 h. Modification of ERα was assessed by Western analysis. (N=3, representative blot shown) **E)** Recruitment of SUMO3 to ERα in a BRET1 assay. HEK293 cells maintained in estrogen-depleted media were transiently transfected with expression vectors for ERα-RLucII and SUMO3-YFP for 48 h and treated with estradiol (10 nM) or antiestrogens (100 nM) for 2.5 h. Net BRET values are shown. The dimerization deficient mutant L507R was used as a negative control to evaluate background BRET activity, which is represented by a dashed line. The absence of statistical significance is identified as "ns", and statistically significant differences are represented by asterisks (* $p \leq 0.05$, ** $p \leq 0.01$, *** $p \leq 0.001$). **F)** Correlation between SUMO3 Net BRET values and relative luminescence counts in reporter assays was determined with a Spearman's rank correlation analysis. Correlation coefficient ($r_s$) and p value (P) are indicated. **G)** Quantification of endogenous ERα protein levels in MCF7 cells maintained in estrogen-depleted media for two days and pre-treated with MG132 for 2 h before incubation with antiestrogens (100 nM) for 6 h. Unmodified bands were quantitated by phosphorimager (N=3) **H)** Correlation between SUMO3 Net BRET values and relative luminescence counts in reporter assays was determined with a Spearman's rank correlation analysis as in F. Correlation coefficient ($r_s$) and p value (P) are indicated.

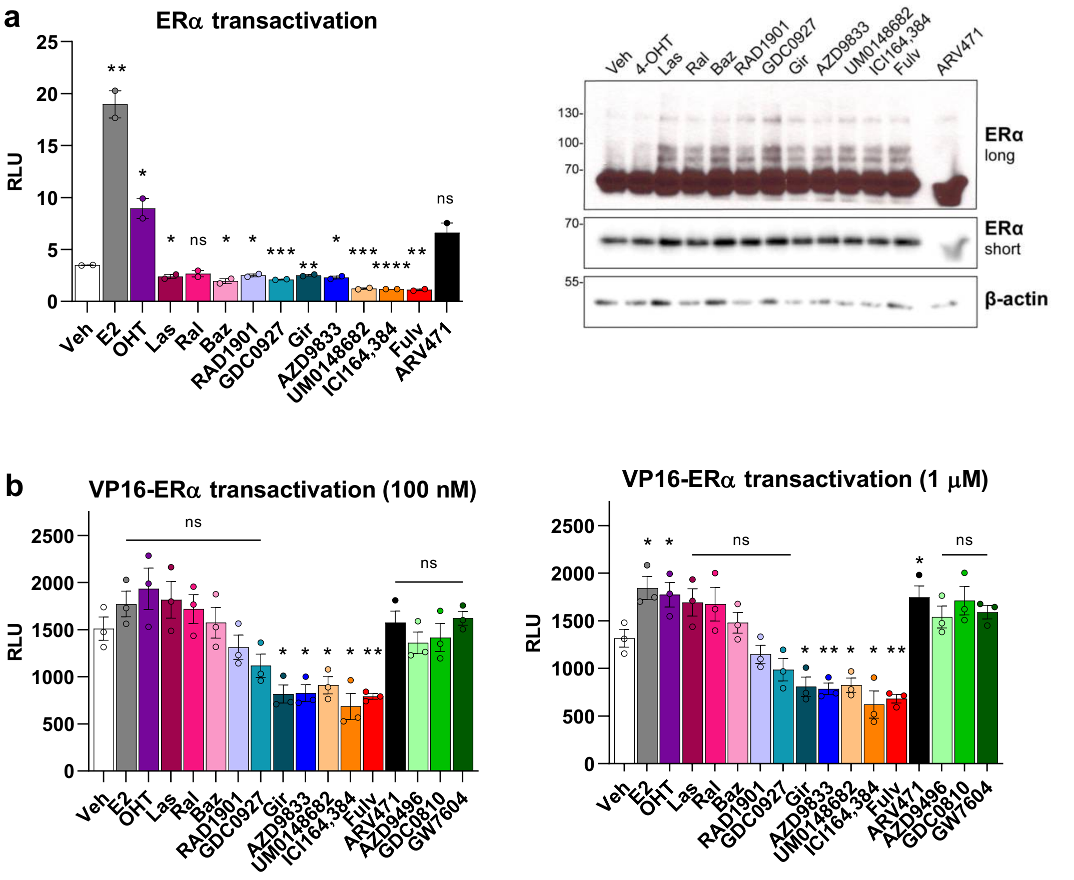


**Supplementary Fig. 3: Induction of SUMOylation correlates with transcriptional repression by ERα even in activated fusion proteins A)** ERα transactivation profile in U2OS transfected cells treated with 100 nM SERMs, SERDs, PROTAC or 10 nM 17β-estradiol (E2). **B)** Effect of different classes of antiestrogens (100 nM) on ERα post-translational modifications in a Western analysis. Western blots show one representative blot out of 2 biological replicates performed in U2OS transiently transfected cells. **C-D)** Transcriptional activity of ERα fused to VP16 activation domain in the presence of ligands. VP16-ERα fusion was transiently transfected in HEK293 cells treated for 1h with antiestrogens at 100 nM, E2 at 10 nM (C) or antiestrogens at 1 μM, E2 100 nM (D). All experiments show mean values of at least 2 biological replicates each performed in triplicate and error bars correspond to standard error of the mean, and error bars correspond to standard error of the mean. Statistical analyses were performed in GraphPad Prism using the Holm-Sidak multiple t-test comparing vehicle and other treatments. The absence of statistical significance is identified as “ns” for p value > 0,05, and statistically significant differences are represented by asterisks (* for p value ≤ 0.05 and ** for p value ≤ 0,01).

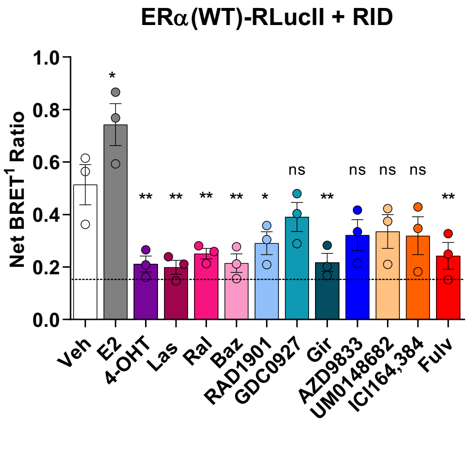


**Supplementary Fig. 4: Impact of antiestrogens on recruitment of AF2 coactivator interacting domain (RID) in a BRET assay.** HEK293 cells maintained in estrogen-depleted media were transiently transfected with expression vectors for ERα-RLucII and RID-YFP for 48 h and treated with estradiol (10 nM) or antiestrogens (100 nM) for 45 min. The dimerization deficient mutant L507R was used as a negative control to evaluate background BRET activity that is represented by the dashed line. The graph shows mean values of 3 biological replicates, each performed with 3 technical replicates. Error bars are standard error of the mean. Statistical analyses were performed in GraphPad Prism using the Holm-Sidak multiple t-test comparing vehicle (Veh) and treatments (“ns” p value > 0.05; * p value ≤ 0.05; ** p value ≤ 0.01).

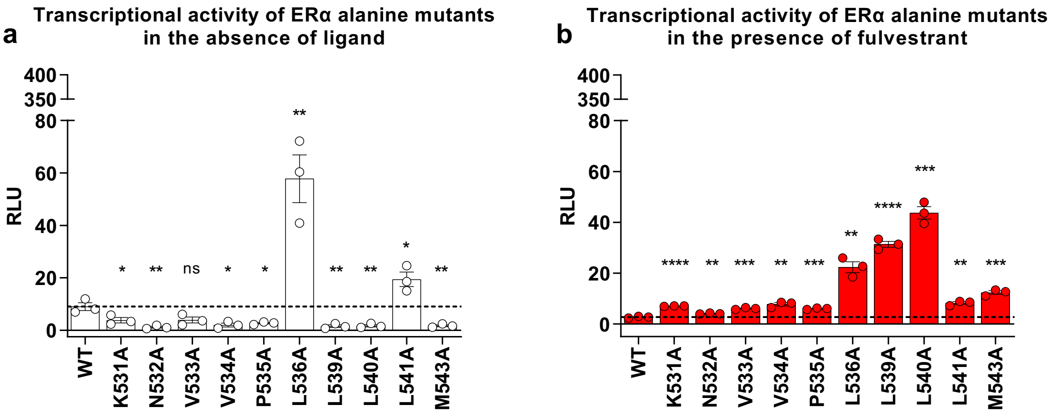


**Supplementary Fig. 5: Contribution of several residues located in H12 for fulvestrant-induced transcriptional repression of ERα in U2OS cells.** A) Transcriptional activity of ERα alanine mutants in the absence (in white) or B) in the presence of fulvestrant (Fulv, in red). U2OS cells were transiently transfected with expression vectors for ERα and GFP and a ERE3-TATA-LUC reporter vector. Cells maintained in estrogen-depleted medium were treated 24h after transfection with 100 nM Fulv for 24 h. Luciferase reporter assays represent mean values of at least 3 biological replicates each performed in triplicates and error bars represent standard error of the mean. Statistical analyses were performed in GraphPad Prism using the Holm-Sidak multiple t-test comparing WT and mutants in the presence or absence of fulvestrant, Absence of statistical significance is identified as "ns" for p value > 0,05, and statistically significant differences are represented by asterisks (* for p value ≤ 0.05, ** for p value ≤ 0,01, *** for p value ≤ 0,001 and **** for p-value ≤ 0,0001).

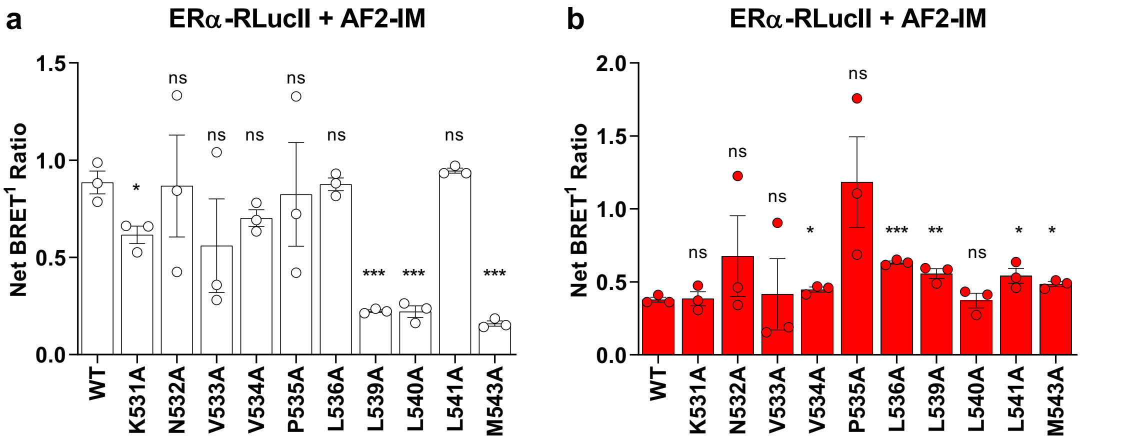


**Supplementary Fig. 6: Impact of point mutations in H12 in the recruitment of AF2 coactivator interacting motif (AF2-IM) to ERα in HEK293. A-B)** Recruitment of AF2 interacting motif (AF2-IM) to ERα alanine mutants in the absence (A) or in the presence (B) of fulvestrant (Fulv). HEK293 cells were transiently transfected with expression vectors for ERα-RLucII and AF2IM-YFP for 48 h and treated with vehicle (A) or fulvestrant 100 nM (B) for 45 min. BRET assays represent mean values of 3 biological replicates each performed in triplicates and error bars represent standard error of the mean. Statistical analyses were performed in GraphPad Prism using the Holm-Sidak multiple t-test comparing WT and mutants in the presence or absence of fulvestrant ("ns" p value > 0.05; * for p value ≤ 0.05; ** p value ≤ 0.01; *** p value ≤ 0.001).

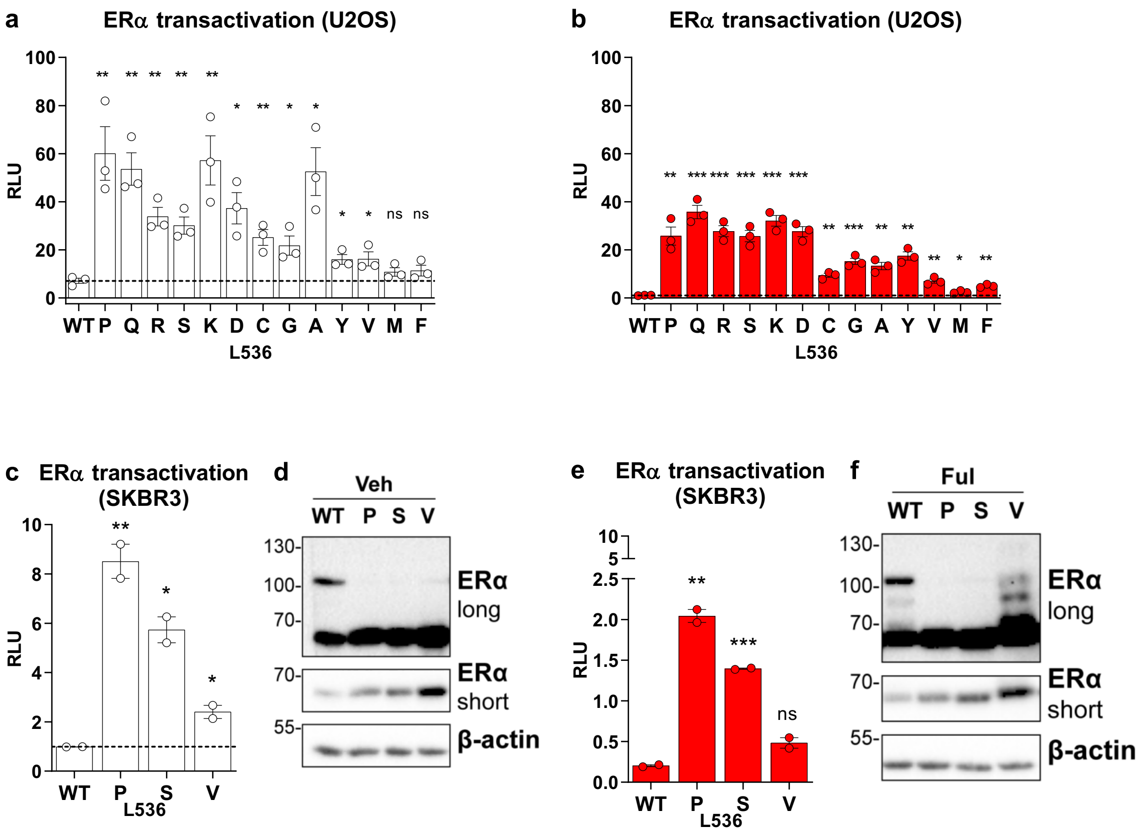


**Supplementary Fig. 7: Contribution of residues at position 536 in fulvestrant-induced transrepression of ERα in ER-negative U2OS and SKBR3 cells. A-B)** Transcriptional activity of ERα L536 mutants in the absence (A) or in the presence of Fulv (B) in U2OS transfected cells. Cells were transiently transfected with expression vectors for ERα, GFP and a ERE3-TATA-LUC reporter vector. Cells maintained in estrogen-depleted medium were treated 24h after transfection with 100 nM Fulv for 24 h. Luciferase reporter assays represent mean values of at least 2 biological replicates each performed in triplicates and error bars represent standard error of the mean. Statistical analyses were performed in GraphPad Prism using the Holm-Sidak multiple t-test comparing WT and mutants in the presence or absence of fulvestrant ( "ns" p value > 0.05; * p value ≤ 0.05; ** p value ≤ 0.01; *** p value ≤ 0.001; **** p-value ≤ 0.0001). **C-D)** Similar experiment performed in ER- breast cancer cells SKBR3. Western analysis was performed on the protein extracts from cells treated for only 1h with ligands.

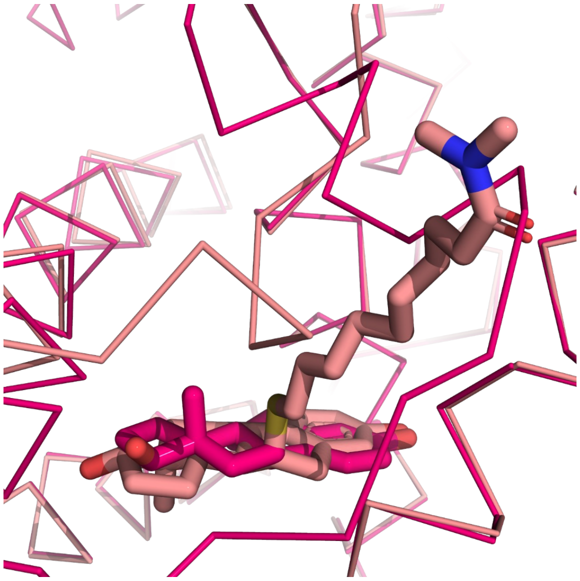

**Supplementary Fig. 8: Superposition of estradiol (fuchsia) and C10(13) (beige) in the ligand binding pocket based on alpha carbon positions.** PDBs: 1GWR and 7N9M

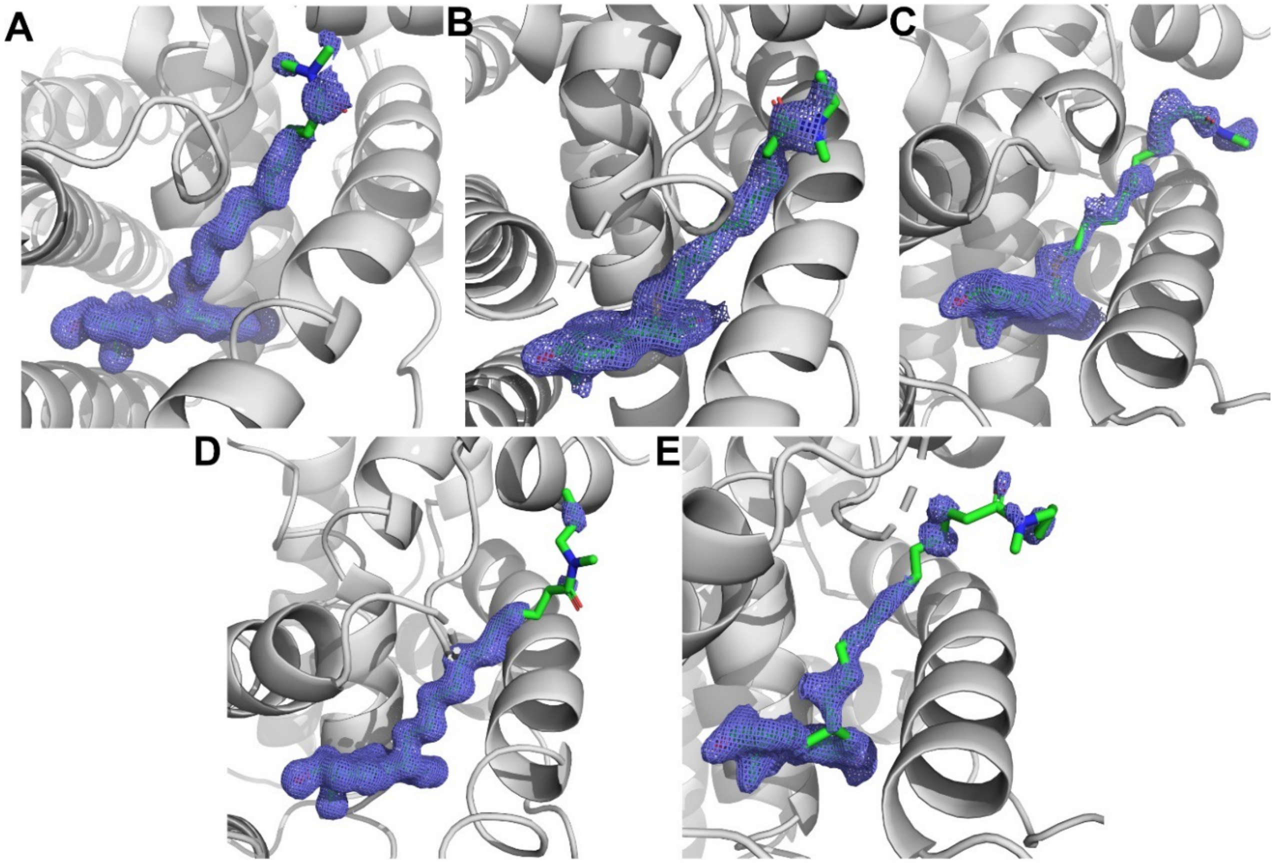


**Supplementary Fig. 9 : 2mFo-DFc difference maps of $C_{10}$(13) (A), $C_{10}$(14) (B), $C_{10}$(15) (C), desmethyl ICI164,384 (D), and ICI164,384 (E) in the ERα ligand binding pocket contoured to 1.0 σ.**

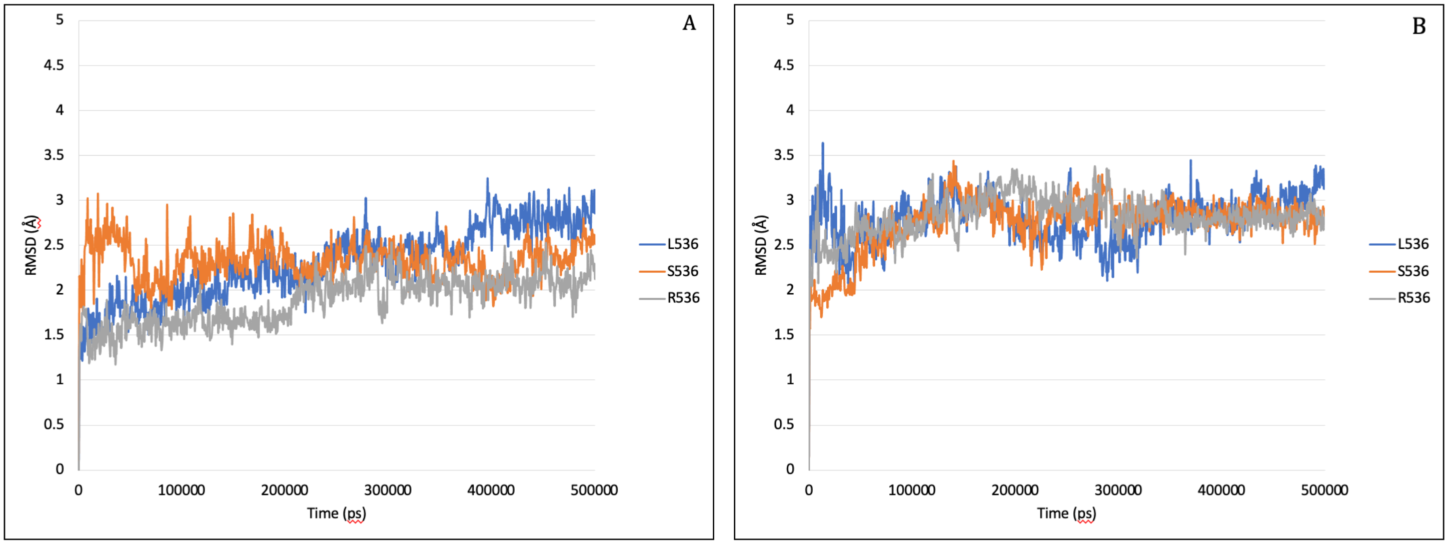


**Supplementary Fig. 10: Root mean square deviation data (RMSD) for the human estrogen receptor alpha LBD structure with different ligands and introduced mutations. A-B** LBD of ERa S536 (orange) L536 (blue) and R536 (grey) with desmethyl ICI164,384 (A) or fulvestrant (B). SD was plotted for the entire protein backbone during MD simulations for 500 ns.

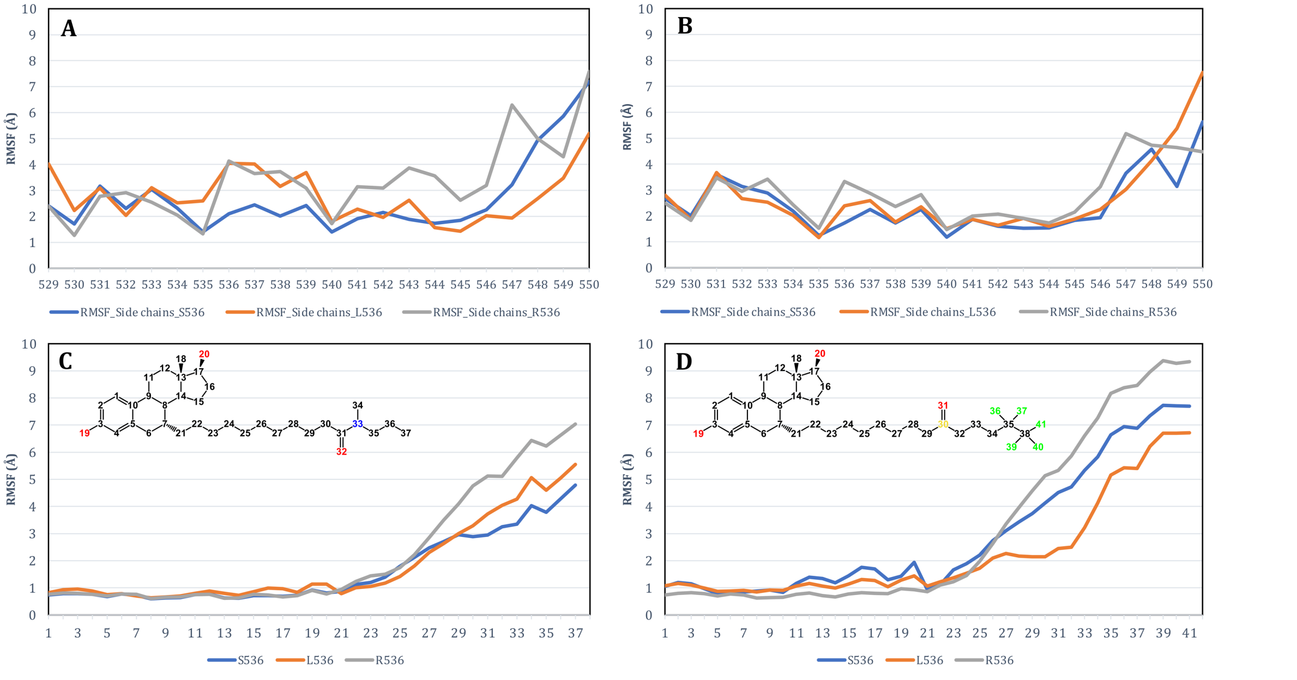


**Supplementary Fig. 11: Root mean square fluctuation data for the side chains of the H11-H12 loop and H12 helix region in all systems over the 500 ns simulation period. A-B** RMSF values are shown for desmethyl-ICI164,384 [A] and for fulvestrant [B] bound to the ERα LBD with S536 (blue), L536 (orange) or R536 (grey). **C-D** RMSF values are shown for residues 529-550 of the ERα LBD with S536 (blue), L536 (orange) or R536 (grey) in the presence of desmethyl ICI164,384 [C] or fulvestrant [D].

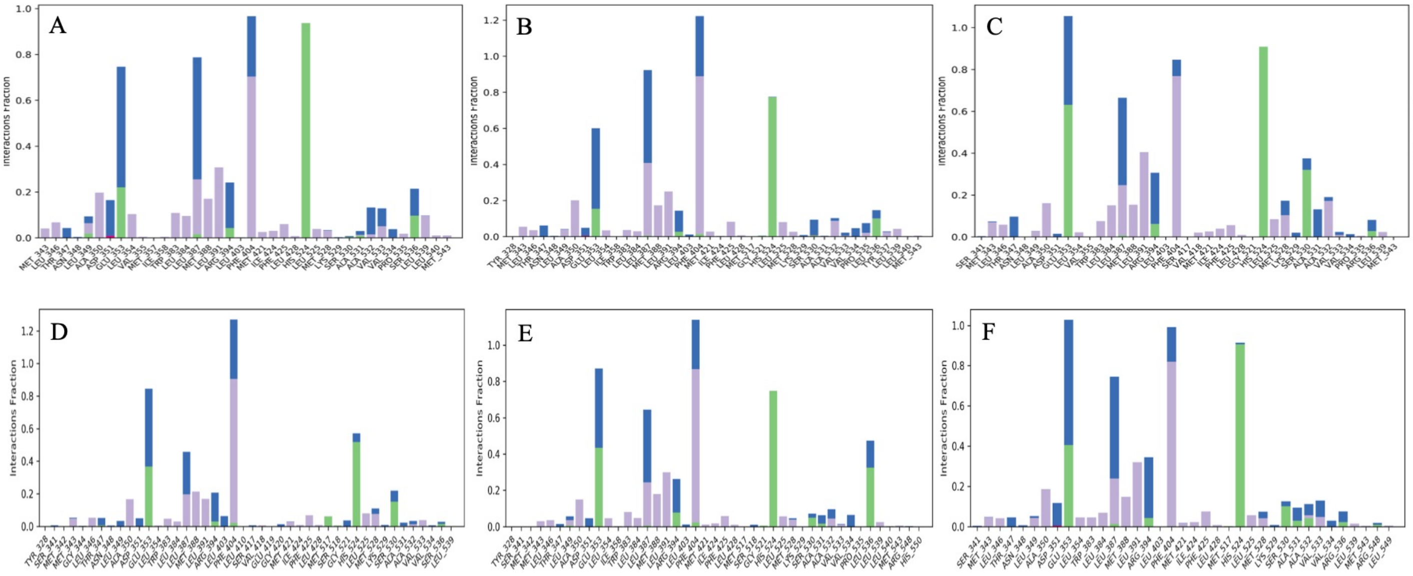


**Supplementary Fig. 12: Interaction of ligands with residues of the ERα ligand binding domain during simulation.** The protein-ligand interaction plot shows the residue list and type of interaction formed for ligands desmethyl ICI164,384 and fulvestrant during the 500 ns simulation time. [A-C] desmethyl ICI164,384 in the L536 [A] S536 [B] or R536 LBD [D-F] fulvestrant in the L536 [D], S536 [E] or R536 LBD [F]. Color code: green: hydrogen bonds – blue: water mediated contact – purple: Van der Waals interactions.
